# Weaker selection on genes with treatment-specific expression consistent with a limit on plasticity evolution in *Arabidopsis thaliana*

**DOI:** 10.1101/2022.10.26.513896

**Authors:** Miles Roberts, Emily B Josephs

## Abstract

Differential gene expression between environments often underlies phenotypic plasticity. However, environment-specific expression patterns are hypothesized to relax selection on genes, and thus limit plasticity evolution. We collated over 27 terabases of RNA-sequencing data on *Arabidopsis thaliana* from over 300 peer-reviewed studies and 200 treatment conditions to investigate this hypothesis. Consistent with relaxed selection, genes with more treatment-specific expression have higher levels of nucleotide diversity and divergence at nonsynonymous sites but lack stronger signals of positive selection. This result persisted even after controlling for expression level, gene length, GC content, the tissue specificity of expression, and technical variation between studies. Overall, our investigation supports the existence of a hypothesized trade-off between the environment specificity of a gene’s expression and the strength of selection on said gene in *A. thaliana*. Future studies should leverage multiple genome-scale datasets to tease apart the contributions of many variables in limiting plasticity evolution.

## 1 Introduction

Organisms must cope with ever-changing environmental conditions to survive and reproduce. If these changes in condition cannot be avoided or escaped, phenotypes that respond to environmental variation through phenotypic plasticity may be adaptive. For example, under low light, the same *Arabidopsis thaliana* genotype will produce more or larger leaves to capture more energy for photosynthesis [64]. Plastic responses are partly controlled through differential gene expression between environments [67, 68]. Understanding the evolution of these condition-specific expression patterns could help reconcile the diversity of plastic responses observed in nature and engineer organisms to overcome environmental challenges.

However, not all organisms can respond plastically to environmental change, so it is crucial to understand the processes that constrain plasticity [80]. These constraints are usually characterized as either costs, where plasticity reduces fitness in some way, or limits to the evolution or maintenance of plasticity [20]. Decades of research has attempted to measure the costs associated with plasticity (reviewed in [69]) but studies often fail to detect costs or find costs that are weak or restricted to certain environments [80, 78, 5]. Theory also predicts that there will be strong selection to alleviate costs [54]. Thus, limits may be more important than costs in shaping the evolution of plasticity.

Recent work suggests that relaxed selection can limit plasticity evolution [71, 54]. For instance, one hypothesis posits that genes are often under selection for environment-specific expression to minimize deleterious pleiotropy [71, 50, 31]. However, narrowing the range of environments where a gene is expressed also reduces the opportunity for negative selection to act on deleterious mutations in the gene [36, 83, 79]. The accumulation of deleterious mutations could then cancel out any selective benefits of the environment-specific expression pattern. Thus, a trade-off arises between a gene’s degree of environment-specific expression and the strength of negative selection acting on said gene. If we assume that environment-specific expression generally contributes to phenotypic plasticity, then this trade-off would potentially limit the maintenance of plasticity [36, 71]. Whether such a trade-off exists has not yet been tested, but the deposition of expression data from hundreds of experimental treatments across hundreds of labs into public repositories now enables approximating environment specificity as treatment specificity and linking treatment-specific expression to the rate of evolution.

One challenge in studying the relationship between treatment specificity and protein evolution is that many factors influence evolutionary rates (for review, see [66, 27, 38, 93]) and these factors are hard to disentangle. A protein’s expression level is often considered the best predictor of its evolutionary rate [66] - a result observed across all domains of life [93] and sometimes considered a “law” of genome evolution [38]. Among multicellular organisms, the degree of tissue specificity in expression is also generally predictive of evolutionary rates [22, 43, 84, 94, 70, 7, 53, 29, 30]. Additional factors that also influence evolutionary rates include exon edge conservation [7], mutational bias [81, 59], gene length [53], gene age [52], GC content [95, 53], expression stochasticity [29], involvement in general vs specialized metabolism [53], identity as a regulatory or structural gene [82], recombination rate [42], codon-bias [6], mating system [85, 28, 62], gene compactness [43, 53], co-expression or protein-protein interaction network connectivity [3, 49, 55, 4, 34], gene body methylation [75], metabolic flux [14], protein structure [47], essentiallity [57, 89, 19], and even plant height [41]. This overabundance of possible explanatory variables suggests that massive genome-scale datasets and careful statistical analysis are required to tease out the influence of treatment-specific expression on evolutionary rates.

To investigate the influence of treatment-specific expression on evolutionary rates, we compiled a dataset of gene expression data across over 200 treatments from over 300 peer-reviewed studies in *A. thaliana*. We annotated RNA-sequencing runs from these studies using standardized ontologies, then processed all of them with the same pipeline. Finally, we combined the resulting gene expression matrix with estimates of selection based on within-species polymorphism and between-species divergence to investigate whether genes with treatment-specific expression were under weaker negative selection.

## 2 Materials and methods

### 2.1 RNA-seq run annotation

We amassed an initial set of RNA-seq runs from the Sustech Arabidopsis RNA-seq database V2 [92] (http://ipf.sustech.edu.cn/pub/athrdb/) excluding any samples not associated with a publication or lacking a tissue type label. On May 24th, 2022 we also downloaded all run metadata from the Sequence Read Archive (SRA) returned by the following search term: (“Arabidopsis thaliana”[Organism] AND “RNA”[Source]) OR (“Arabidopsis thaliana”[Organism] AND “RNA-Seq”[Strategy]) OR (“Arabidopsis thaliana”[Organism] AND “TRANSCRIPTOMIC”[Source]). All SRA runs were linked to their associated publications, if possible, using Entrez. Any SRA run numbers that we could not link to a PUBMED ID or DOI were omitted. We then manually removed all SRA runs that originated from transgenic, mutant, hybrid, grafted, cell culture, polyploid, or aneuploid samples based on information in the SRA metadata and associated publications. Runs from any naturally-occurring *A. thaliana* accession were included. We also omitted SRA runs that focused on sequencing non-coding RNA (ncRNA-seq, miRNA-seq, lncRNA-seq, sRNA-seq, etc.). After applying these criteria, any bioprojects with 8 or fewer SRA run numbers remaining were also omitted.

All runs were labeled with treatment and tissue type descriptions using the Plant Experimental Conditions Ontology (PECO) and the Plant Ontology (PO) [15], respectively, based on information in their associated publications and SRA metadata. In our analysis, control exposure was defined as long day conditions (12 hrs light exposure or longer, but not constant light) and growing temperatures in the range of 18° - 26°, inclusive, without explicit application of stress or nutrient limitation. Warm treatments were defined as 27° or higher, while cold treatments were defined as 17° or lower. Any studies that did not report both day length and growing temperature were omitted. Any runs that could not be linked to treatments based on their annotations in the SRA or Sustech databases were also omitted. Treatment with polyethylene glycol (PEG) was categorized as drought exposure. Samples from plants that were recovering from stress were categorized according to the growth conditions of the recovery state instead of the stressed state. When appropriate, we labeled samples with multiple PECO terms. For example, a sample that was subjected to both heat stress and high light stress would get two PECO terms (one for each stress) and be treated separately from samples subjected to only heat stress or only light stress. Tissue type labels were eventually collapsed to the following categories: whole plant, shoot, root, leaf, seed, and a combined category of flower and fruit tissues. The flower and fruit tissue categories were combined because of their developmental relationship and small size relative to the other categories. In the end, we had a dataset of 24,101 sequencing runs from 306 published studies.

### 2.2 RNA-seq run processing

All RNA-seq runs were processed using the same workflow to remove the effects of bioinformatic processing differences between studies on expression level. First, runs were downloaded using the SRA toolkit (v2.10.7), but 90 runs were not publicly available and thus failed to download. All successfully downloaded runs were trimmed using fastp v0.23.1 [10], requiring a minimum quality score of 20 and a minimum read length of at least 25 bp (-q 20 -l 25). Trimming results were compiled using multiqc v1.7 [25]. All trimmed runs were then aligned to a decoy-aware transcriptome index made by combining the primary transcripts of the Araport11 genome annotation [11] with the *A. thaliana* genome in salmon v1.2.1 [61] using an index size of 25bp. The salmon outputs of each run were then combined with a custom R script to create an gene-by-run expression matrix. We omitted 423 runs with a mapping rate ¡ 1 %, 215 runs with zero mapped transcripts, and 18 genes with zero mapped transcripts across all runs from further analysis. We note that although this cut-off does not exclude samples with more modest mapping rates (e.g. 20 - 60 %) the choice to include these samples was to avoid removing large chunks of data as “outliers” and analyzing only those samples that conform to our expectations.

### 2.3 Whole genome sequence data processing

We downloaded whole genome sequencing data for 1135 *A. thaliana* accessions from the 1001 genomes project panel (SRA project SRP056687) [2] using the SRA toolkit. All runs were trimmed using fastp [10], requiring a minimum quality score of 20 and a read length of at least 30 bp (-q 20 -l 30). Trimmed reads were then aligned to the *A. thaliana* reference genome using BWA v0.7.17 [46]. The alignments were sorted and converted to BAM format with SAMTOOLS v1.11 [18], then optical duplicates were marked with picardtools v2.22.1. Haplotypes were called for each accession, then combined and jointly genotyped with GATK v4.1.4.1 assuming a sample ploidy of 2, heterozygosity of 0.001, indel-heterozyogsity of 0.001, and minimum base quality score of 20. Invariant sites were included in the genotype calls with the –include-non-variant-sites option. All calls were restricted to only coding sequence (CDS) regions based on the Araport11 annotation by supplying a BED file of CDS coordinates made with bedtools (v2.29.2). Following [39], variant and invariant sites were filtered separately using both GATK and vcftools v0.1.15 [17]. Variant sites were filtered if they met any of the following criteria: QD *<* 2, QUAL *<* 30, MQ *<* 40, FS *>* 60, HaplotypeScore *>* 13, MQRankSum *<* -12.5, ReadPosRankSum *<* -8.0, mean depth *<* 10, mean depth *>* 75, missing genotype calls *>* 20%, being an indel, or having more than 2 alleles. In the end, 1,915,859 variant sites across all coding sequences were retained for further analysis. Invariant sites were filtered if they met any of the following criteria: QUAL *>* 100, mean depth *<* 10, mean depth *>* 75, missing genotype calls *>* 20%. Finally, variant sites were annotated using snpEff (Java v15.0.2) [13] and variants labeled as either missense or synonymous were separated into different files using SnpSift [12].

### 2.4 Selection estimated from between-species divergence

We identified 1:1 orthologs between the primary transcripts of *A. thaliana* and *Arabidopsis lyrata* with Orthofinder v2.5.4 [24]. For each 1:1 ortholog, we aligned their protein sequences with MAFFT L-INS-I v7.475 [35], then converted the protein alignments to gapless codon-based alignments using pal2nal v14 [73]. Using the gapless codon-based alignments, we estimated *dN/dS* using the method in [56] implemented as a custom Biopython v1.79 script and implemented through the codeml program in the PAML package v4.9 [91]. Unlike codeml, the custom Biopython script also returns counts of nonsynonymous (*N*) and synonymous sites (*S*) within each gene as described in [56], which we later used to calculate nucleotide diversity per nonsynonymous site (*π*_*N*_) and per synonymous site (*π*_*S*_). Before proceeding with more analyses, we confirmed that our estimates of *dN* and *dS* were consistent between our Biopython script and codeml (Figure S5, Pearson correlations *dN* : *ρ* = 0.9998, *dS* : *ρ* = 0.9809). The outputs of the Biopython script were used in all subsequent analyses.

### 2.5 Selection estimated from within-species polymorphism

#### 2.5.1 Nucleotide diversity at nonsynonymous sites

Nucleotide diversity (*π*) was calculated for each gene with pixy v1.2.3.beta1 [39] three times: once using all sites (both variant and invariant), once using missense sites plus invariant sites, and once using synonymous sites plus invariant sites. These estimates were then converted to *π, π*_*N*_, and *π*_*S*_, respectively, by first multiplying the per site estimate output from pixy by the number of sites included in the analysis. Then, to get *π*_*N*_ and *π*_*S*_, the values from analyses of missense plus invariant, and synonymous plus invariant sites were divided by the *N* and *S* values for each gene, respectively, as determined by the method in [56].

#### 2.5.2 Tajima’s D

We next calculated Tajima’s D for each gene. First, we calculated *π* and Watterson’s Theta (*θ*_*W*_) for each variant site *i* within a gene (*π*_*i*_ and *θ*_*W i*_ respectively). In this case, *π*_*i*_ was calculated as:

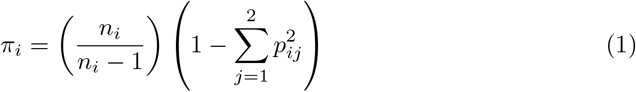

Where *n*_*i*_ is the number of sequenced chromosomes with non-missing genotypes for variant *i, p*_*i*1_ is the frequency of the reference allele, and *p*_*i*2_ is the frequency of the alternative allele. Then, *θ*_*W i*_ was calculated as:

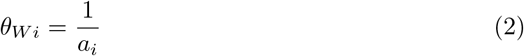

Where *a*_*i*_ is:

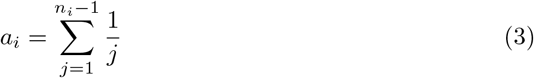

This calculation of *θ*_*W i*_ is equivalent to the usual calculation of *θ*_*W*_ with the number of segregating sites set to one. Next, the variance in Tajima’s D was calculated for each site as:

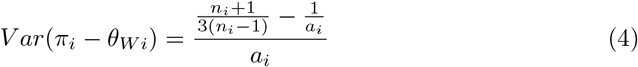

This is equivalent to equation 38 in [74] with the number of segregating sites set to one. Finally, the results of the above calculations were combined in the following formula:

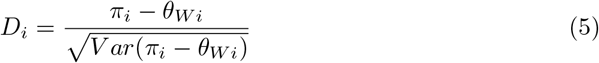

To get Tajima’s D for each gene, we then averaged across the *D*_*i*_ values for all the variant sites within a gene.

#### 2.5.3 Direction of Selection (DoS)

Counts of nonsynonymous and synonymous polymorphisms within each gene (*P*_*N*_ and *P*_*S*_, respectively) were determined with bedtools (v2.29.2). The number of nonsynonymous and synonymous differences (*D*_*N*_ and *D*_*S*_, respectively) between *A. thaliana* genes and their 1:1 *A. lyrata* orthologs, if present, were estimated during the process of calculating *dN/dS* in Biopython as described above. These values were then used to calculate the direction of selection (DoS) [72] as follows:

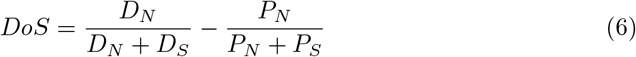

We chose this metric, as opposed to the proportion of amino acid substitutions driven by positive selection (*α*), because it is less biased than *α* [72] and was successfully used in studies similar to ours [60]. Furthermore, we found that *α* often returns uninterpretable negative values when applied to *A. thaliana*, perhaps because of an excess of slightly deleterious polymorphisms [58] due to their predominantly selfing mating system [9].

### 2.6 Treatment specificity

Treatment specificity (*τ*) was estimated separately for runs from each tissue type using the following formula [88]:

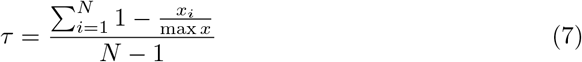

Where *x* is the vector of average expression values of a gene in each treatment category, measured in transcripts per million (TPM), and where *N* is the number of treatment categories. Dividing by *N* means that *τ* varies between zero and one, where zero indicates no specificity and one indicates exclusive specificity to a single treatment. We used this metric of specificity because it is consistently more robust than others [40] and is normalized by the number of treatments included, making it comparable across data sets. We also applied the same formula to calculate tissue specificity in several different treatment conditions.

### 2.7 Simulating correlations between average expression and specificity index

Average expression level and measures of expression specificity are correlated by definition because genes with more treatment/tissue-specific expression will have lower average expression across all treatment/tissue categories. We ran two simulations to better illustrate the factors driving the correlation between average expression and the specificity index, *τ*. In both simulations, we generated 1000 random matrices, where each element *x*_*ij*_ represented the expression of gene *i* in experiment *j*, by sampling from a zero-inflated negative binomial distribution:

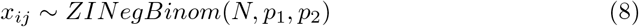

Where the size and probability parameters of the negative binomial component were *N* = 100 and *p*_1_ = 0.1, respectively, while the probability of an expression value being non-zero was *p*_2_ = 0.4. All matrices included 5 groups of columns, with 5 columns per group, representing replicates of tissue/treatment groups. For both simulations, we averaged across columns within each group to simulate the calculation of tissue/treatment-wide averages. We then applied the formula for *τ* across the rows of this averaged matrix to get expression specificity. In one simulation, we calculated expression level by averaging across the rows of the expression matrix. In a second simulation, we excluded experiments where a gene was not expressed (*x*_*ij*_ = 0) from the calculation of average expression.

### 2.8 Average expression, length, GC content, family size

Calculating the average expression of each gene was a three-step process. First, we averaged together runs with matching SRA experiment IDs because these runs represented technical replicates of the same biological sample and treatment conditions. Second, we partitioned our gene-by-experiment expression matrix by the tissue type each sample came from. Finally, for each tissue type’s expression matrix, we averaged across all of the expression values of each gene across all experiments, excluding values ¡ 5 transcripts per million (TPM). We excluded values ¡ 5 TPM from the average expression calculation to avoid a high correlation between average expression and treatment-specificity, as has been reported in previous studies [70]. This high correlation occurs because an environment-specific gene will by definition also have low average expression across environments it is rarely expressed in. Furthermore, we excluded values ¡ 5 TPM to avoid including small expression values that could be artifacts of alignment error.

The length and GC content of each gene was measured using the bedtools nuc command (v2.29.2) and included each gene’s introns and untranslated regions when present. We included introns and untranslated regions in the estimate of gene length because they play important roles in determining rates of protein evolution [8, 23]. Finally, the family size for each gene was estimated as the number of *A. thaliana* genes in their respective orthogroups output by OrthoFinder.

### 2.9 Partial correlation analysis

Not all treatment-tissue combinations were sampled in the overall RNA-seq dataset, causing confounding between the treatment and tissue labels. We resolved this in two ways. First, we subset the data to only the treatment conditions where all tissue types were represented. Second, we subset the data by tissue type and analyzed each subset separately. For each subset, we calculated partial spearman correlations between treatment specificity and our measures of selection (*dN, π*_*N*_, Tajima’s D, and *DoS*) after accounting for average expression (excluding values TPM ¡ 5), gene length, and GC content using the ppcor R package [37]. For partial correlation analyses involving *π*_*N*_ and Tajima’s D, we also controlled for gene family size. We did not account for gene family size in partial correlation analyses involving *dN* or *DoS* because these metrics apply to only genes with one family member in this study. When calculating partial correlations involving *dN*, we excluded any genes with saturating divergence (*dS >* 1). All statistical analyses and data visualizations used R v4.0.3 and used color palettes in the scico R package [16, 63].

### 2.10 Surrogate variable analysis

We recalculated treatment specificity and repeated all partial correlation analyses after correcting each data subset for technical between-experiment variation (i.e. batch effects), following an approach from [26]. Batch effects include variables that influence gene expression measurements but are not of interest to this study, such as the sequencing platform and the library prep protocol used in each experiment. First, with our data already subset by tissue type, we further subset to only include treatments with RNA-seq runs from at least two studies. This minimizes confounding between-treatment variation with the technical between-experiment variation we aimed to account for. We then applied surrogate variable analysis (SVA) using the svaseq() function within the SVA package [45] to each of these subsets. Briefly, SVA models gene expression as:

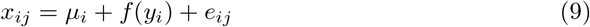

Where *x*_*ij*_ is the expression of gene *i* in experiment *j, μ*_*i*_ is the average expression of gene *i* across all experiments, and *y*_*i*_ is the value of a predictor variable of interest for gene *i*. Furthermore, *f* (*y*_*i*_) gives the deviation of gene *i* from its average expression based on the value of *y*_*i*_ and *e*_*ij*_ is the residual error. SVA takes this model and partitions the residual variance, *e*_*ij*_, into:

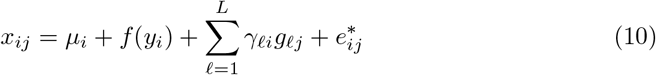

Where 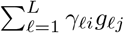 gives the summed effects of *L* unmodeled variables (*g*_*ℓj*_) for each gene and 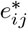 gives the gene-specific noise in expression. SVA does not attempt to estimate what the unmodeled variables influencing expression are, but rather find a set of vectors (the surrogate variables) that span the same space as g:

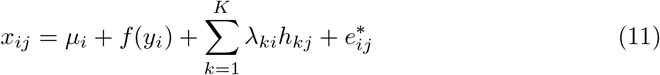

Where each h_k_ is a surrogate variable and each *λ*_**k**_ gives the effects of each surrogate variable on gene expression. For our analyses, our predictor variable *y*_*i*_ was treatment type. To get a measure of expression where the effects of surrogate variables are removed, we then subtracted off the effects of surrogate variables from both sides of the above equation.

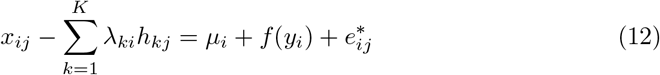

Where 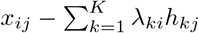 gives us our expression values accounting for the effects of surrogate variables. The net result here is a reduction in the amount of unexplained or seemingly stochastic variation in expression because sources of variation have been attributed to “surrogates” that span the same space as real batch variables. We also conducted principal component analysis in R before and after SVA to verify the removal of batch effects.

## 3 Results

### 3.1 Summary of tissue differentiation, treatment specificity, and selection in overall dataset

To understand how treatment specificity of gene expression affects evolutionary rates of proteins, we queried the Sequence Read Archive for all *A. thaliana* RNA-seq experiments published before May 2022. We then annotated these experiments with standardized tissue and treatment ontology terms, manually filtered the dataset, and then processed all RNA-seq runs with a standardized pipeline. The number of sequencing experiments associated with each combination of tissue and treatment labels is summarized in Tables S1. Overall, the most sampled tissue category was leaf (4,642 experiments) followed by root (3,348 experiments), whole plant (2,492 experiments), seed (1,866 experiments), shoot (1,106 experiments), then fruit and flower (266 experiments). The four most sampled treatment categories were control (5,701 experiments), cold air exposure (675 experiments), short day length (561 experiments), and short day length plus *Botrytis cinerea* exposure (407 experiments). Any sequencing runs that shared an SRA experiment ID were averaged to produce individual gene expression values for each SRA experiment.

We first looked at the distribution of mapping rates across all RNA-seq runs. The median mapping rate was 72.39 % (Figure S1) and we excluded runs with a mapping rate ¡ 1% from further analyses. We next performed a principal components analysis (PCA) on the expression matrix and observed strong differentiation between root and non-root tissues along PC2 (Figure 1). We also observed that nearly all genes had some degree of treatment specificity in their expression (Figures 2A, S3). Furthermore, only a small proportion of genes had strong signatures of selection based on *dN/dS, π*_*N*_ */π*_*S*_, DoS, or Tajima’s D (Figure 2B-D, Figure S2). The treatment specificity of expression was lower on average in flower and fruit tissue compared to the other tissues (Figure S3). However, tissue specificity did not vary widely depending on the treatment condition (Figure S4).

**Figure 1.**
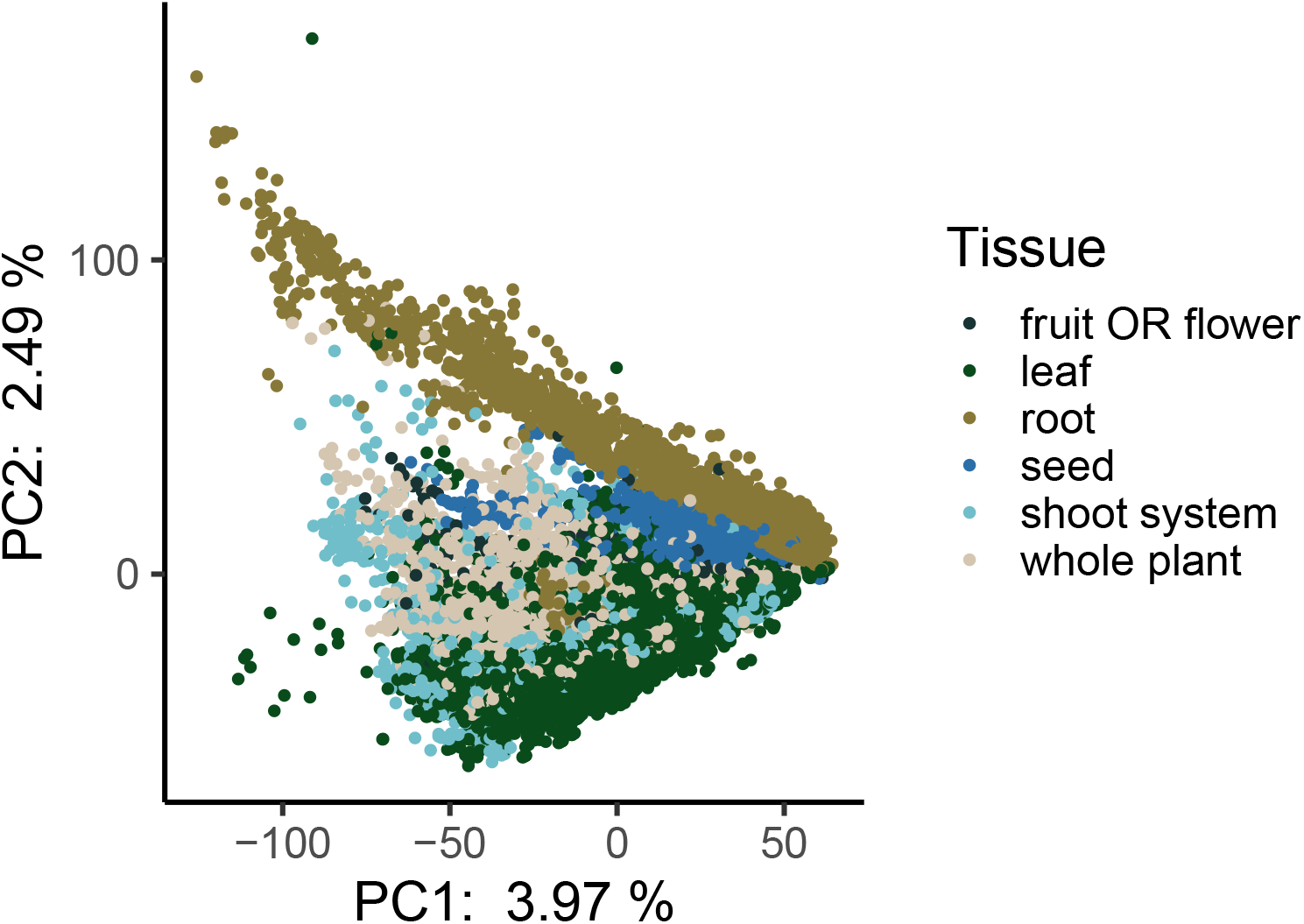
Principal components analysis of all expression data. Each point represents a different RNA-seq experiment and is colored by its associated tissue type. Experiments from all treatment conditions are included in this analysis. Plotting order was randomized to avoid overplotting.

**Figure 2.**
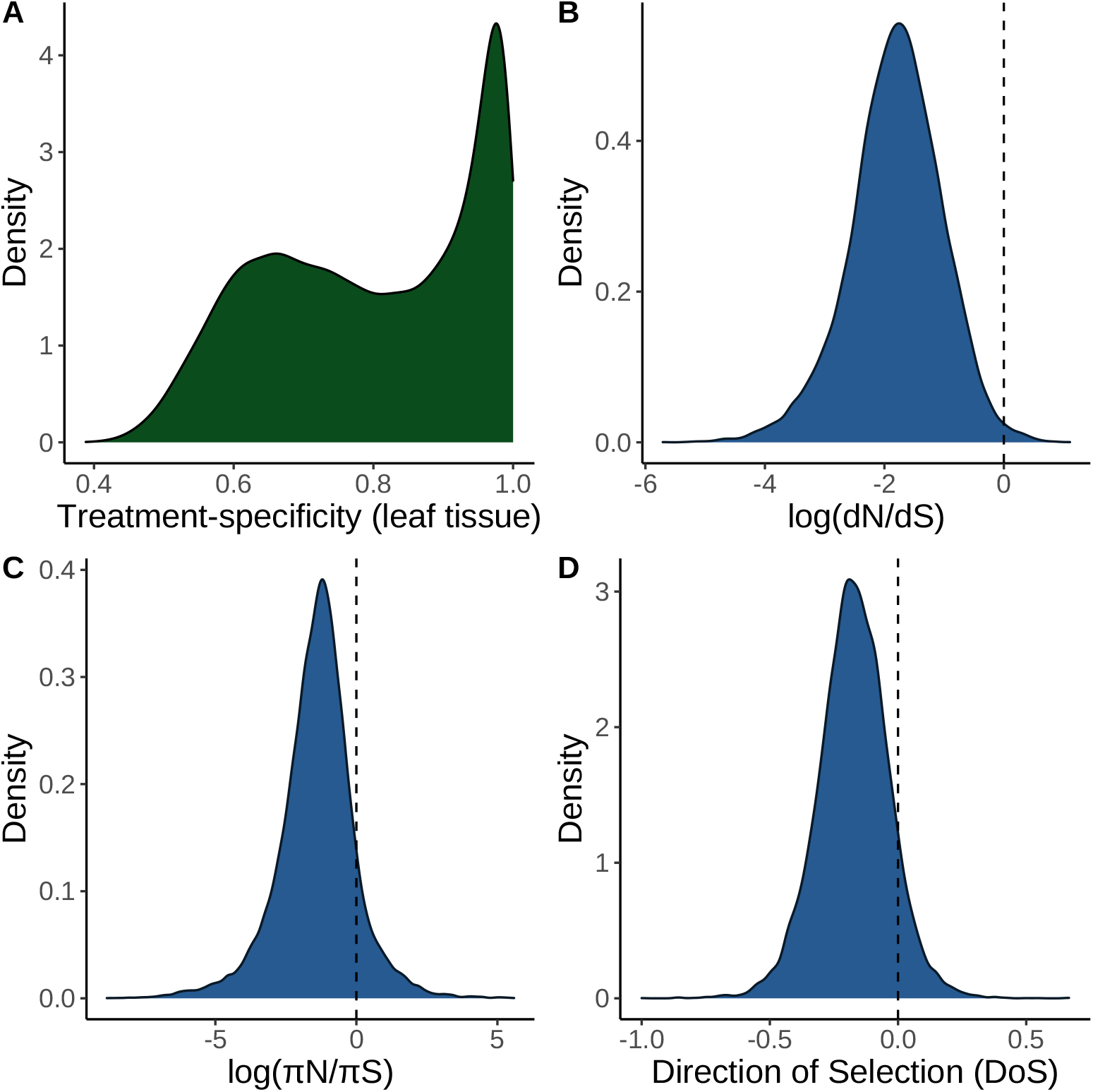
Density plots of key variables measured in this study. (**A**) Distribution of treatment specificity in leaf tissue expression across all genes included in this study. The area underneath the curve in a given interval of treatment specificity represents the proportion of genes in this study that fall within that range of treatment specificity. (**B**) Distribution of *dN/dS* across all genes included in this study. The area to the right of the dashed line represents the proportion of genes in this study with *dN/dS >* 1. (**C**) Distribution of *π*_*N*_ */π*_*S*_ across all genes included in this study. The area to the right of the dashed line represents the proportion of genes in this study with *π*_*N*_ */π*_*S*_ *>* 1. (**D**) Distribution of DoS across all genes in this study. Area to the right of the dashed line represents the propotion of genes with DoS ¿ 0, which is interpreted as evidence of adaptive evolution.

### 3.2 Omitting samples with low expression disentangles expression level and specificity

Genes that are only expressed in one treatment or tissue will, by definition, have low mean expression across all environments or tissues [86]. Thus, we sought a method of calculating expression level that was independent of treatment specificity. To better understand the relationship between average expression and treatment specificity, we calculated correlations between treatment-specificity and expression level while either including or excluding low expression values (TPM ¡ 5) on our real RNA-seq dataset. We found that excluding low expression values decreased the correlation between average expression and treatment-specificity in leaf tissue samples (Figure 3) and other tissues (Figures S34 - S38) and replicated the result by simulating gene expression matrices (Figure S39). Thus, for all later partial correlation analyses (see next section) we quantified each gene’s average expression after dropping experiments where the gene was not expressed (TPM ¡ 5).

**Figure 3.**
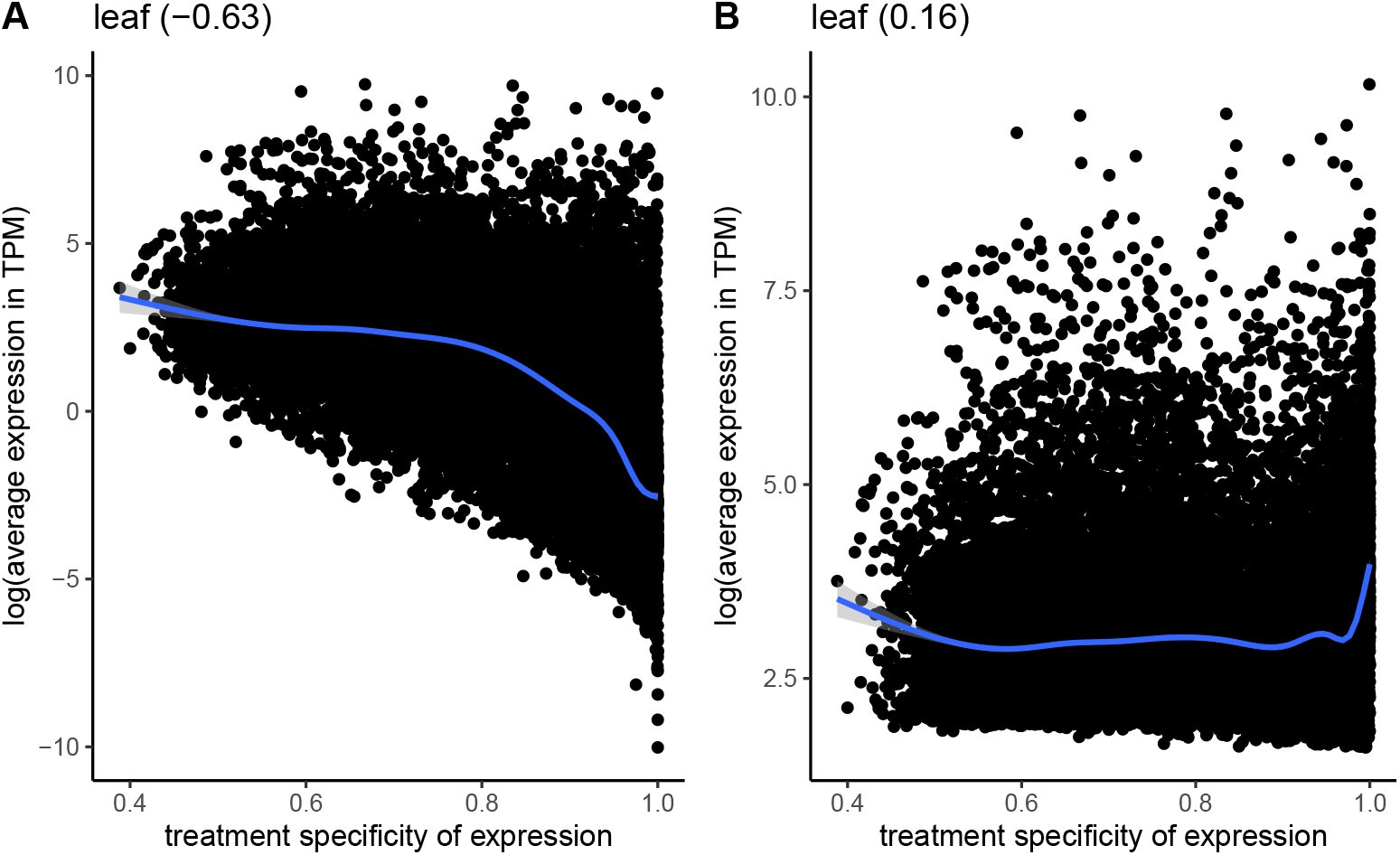
Correlation between the average expression in transcripts per million (TPM) and treatment specificity of genes when samples with low expression (*<* 5 TPM) are included (**A**) vs excluded (**B**). Expression level and treatment specificity were calculated using only data from leaf tissue samples. Line is a smoothing line with 95 % confidence intervals and values in parentheses give spearman correlation.

### 3.3 Treatment specificity correlates with levels of nonsynonymous diversity and divergence in genes

We next calculated partial correlations between treatment specificity and measures of selection after controlling for average expression, gene length, GC content, and tissue specificity in expression. These partial correlations were calculated separately for expression data on each tissue type and did not account for batch effects (see next section). Among leaf tissue samples, average expression had significant partial correlations with *dN* (*ρ* = −0.19, p-value = 2.1 *×* 10^−122^) and *π*_*N*_ (*ρ* = −0.17, p-value = 2.8 *×* 10^−175^) after controlling for other factors (Figures 4A,4B). Treatment specificity was more strongly correlated with *dN* (*ρ* = 0.10, p-value = 7.6 *×* 10^−31^) and *π*_*N*_ (*ρ* = 0.10, p-value = 1.2 *×* 10^−62^) than Tajima’s D (*ρ* = 0.01, p-value = 3.1 *×* 10^−7^) and DoS (*ρ* = 0.04, p-value = 2.3 *×* 10^−06^, Figure 4C, 4D). Furthermore, the top 25% most treatment-specific genes in leaf tissue for our dataset have average *dN* and *π*_*N*_ values nearly 2.5 times greater than the 25% least treatment-specific genes (*dN* = 0.025 vs 0.061; *π*_*N*_ = 0.0014 vs 0.0032). Meanwhile, the most and least treatment-specific genes have average Tajima’s D values of are -0.44 and -0.43, respectively, and average *DoS* values of -0.19 and -0.14, respectively. The strongest partial correlation generally occurred between tissue specificity and treatment specificity (Spearman’s *ρ* = 0.53 − 0.60, Figure 4). Gene family size had among the weakest partial correlations with *π*_*N*_ compared to other covariates, but strongly correlated with treatment specificity (*ρ* = 0.12, p-value = 6.3 *×* 10^−84^, Figure 4B). All of these findings generally held when average expression and treatment specificity were calculated on data from other tissues (Table S2, Figures S6-S10).

**Figure 4.**
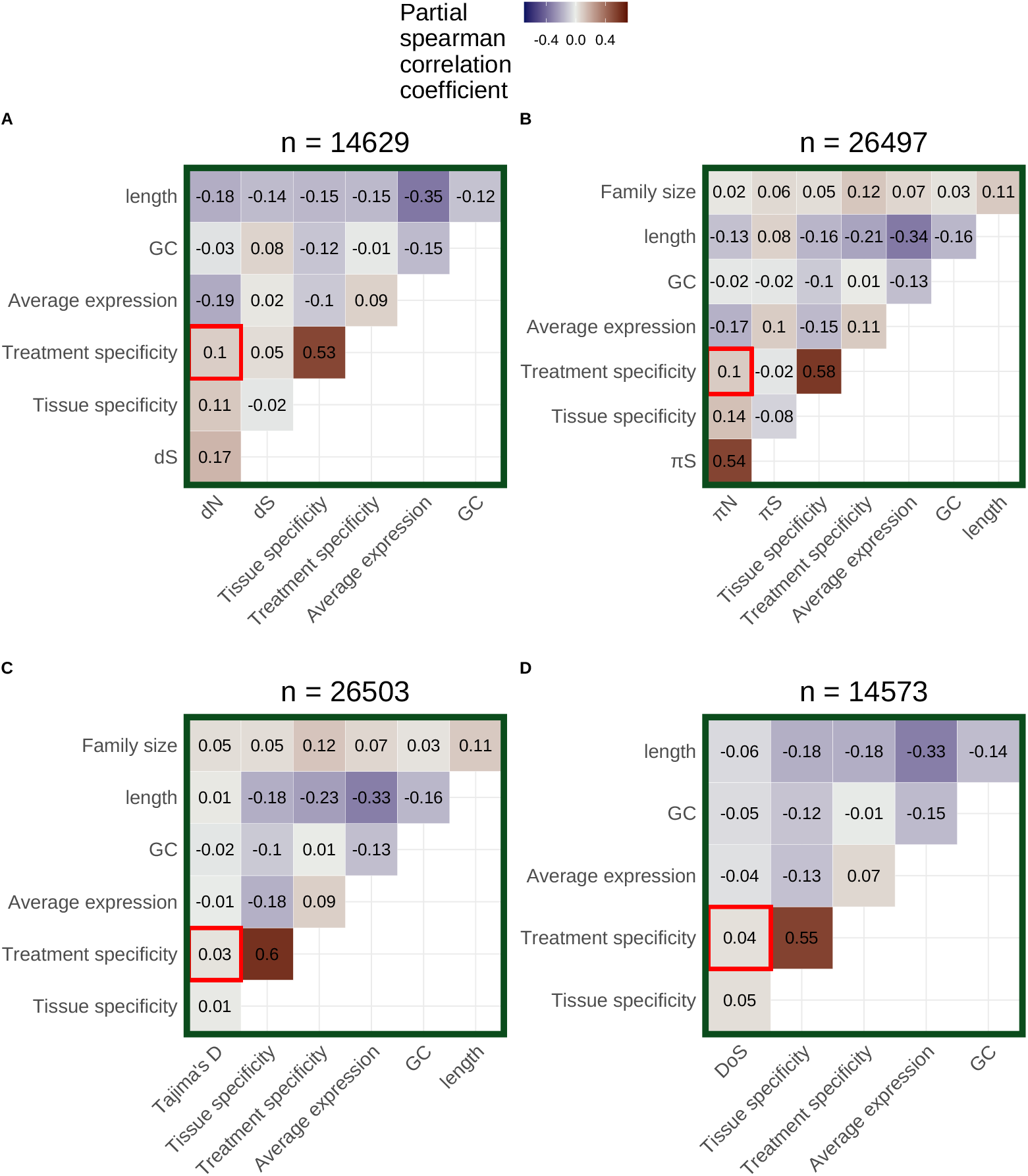
Partial correlation analysis including either (**A**) *dN*, (**B**) *π*_*N*_, (**C**) Tajima’s D, or (**D**) direction of selection (DoS) as a covariate. Average expression excludes values ¡ 5 TPM and was calculated using only leaf tissue samples. Treatment specificity was also calculated using only leaf tissue samples. Tissue specificity was calculated using only control samples across all tissue categories. The number of genes included in each partial correlation analysis (n) is listed at the top of each heatmap.

### 3.4 Correlations between treatment specificity and nonsynonymous variation persist after controlling for batch effects and dataset imbalance

While combining gene expression data across multiple studies can increase the statistical power of an analysis, there are some potential concerns. First, if many tissue-treatment combinations are not sampled, the dataset will be unbalanced and the effects of tissue and treatment variation on expression could be confounded. Consistent with this expectation, there was a high correlation between tissue specificity and treatment specificity in our initial analyses (Figure 4, S6-S10). Furthermore, combining data from multiple laboratories could generate batch effects [44]. To address the issues of imbalance and batch effects, we first subset our data to only include treatments where all tissue types were represented. This subset included the treatments of control, abscisic acid, continuous light, warm/hot air temperature, and cold air temperature. We then used SVA to correct for the influence of unknown batch effects on this data subset [45]. After SVA, treatment specificity positively correlated with *dN* (*ρ* = 0.10, p-value = 1.6 *×* 10^−32^) and *π*_*N*_ (*ρ* = 0.07, p-value = 1.5 *×* 10^−23^) when average expression and treatment specificity were calculated on combined fruit and flower data (Figures S33). However, treatment specificity in other tissue types generally did not correlate with our measures of selection (Figures S28-S33, Table S4).

The inclusion of only five treatments in the above analysis could limit quantification of a gene’s treatment specificity. Thus, in order to include data from a larger number of treatments, avoid dataset imbalance, and avoid batch effects, we split our expression matrix into six subsets by tissue category. We then further removed treatments that only had expression data from one study to avoid confounding treatment effects with study-specific batch effects. We applied SVA [45] to each of these tissue-specific subsets. After SVA, the expression profiles of most genes appear less treatment-specific (Figures S16-S21 panels A vs B). We also observed less separation in PCA space within treatment groups after SVA (for example, see Figures S16C and S16D). Average expression levels before SVA were generally correlated with expression levels after SVA (Figures S16-S21 panels A and B). In partial correlations on each SVA-corrected subset, treatment specificity significantly correlated with *dN* (*ρ* = 0.13, p-value = 6.9 *×* 10^−50^) and *π*_*N*_ (*ρ* = 0.16, p-value = 3.9 *×* 10^−128^) but less strongly correlated with Tajima’s D (*ρ* = 0.04, p-value = 6.6 *×* 10^−10^) and DoS (*ρ* = 0.05, p-value = 2.0 *×* 10^−8^) for the leaf tissue data subset (Table 1, Figures 5). These patterns were similar in other tissue types (Figures S11-S15, Table S3).

**Table 1.**
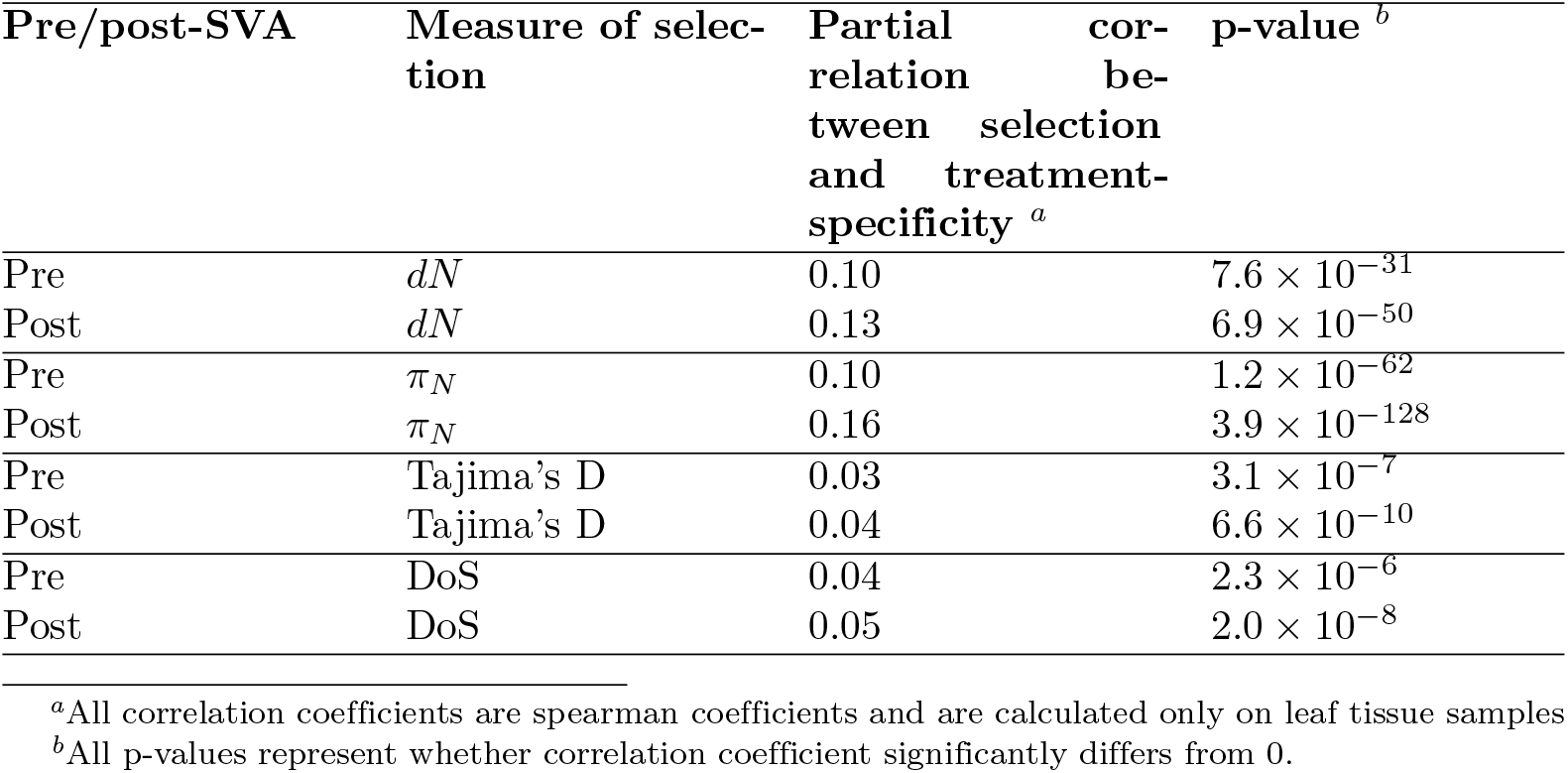
Partial correlations between treatment-specificity and different measures of selection pre-SVA and post-SVA

**Figure 5.**
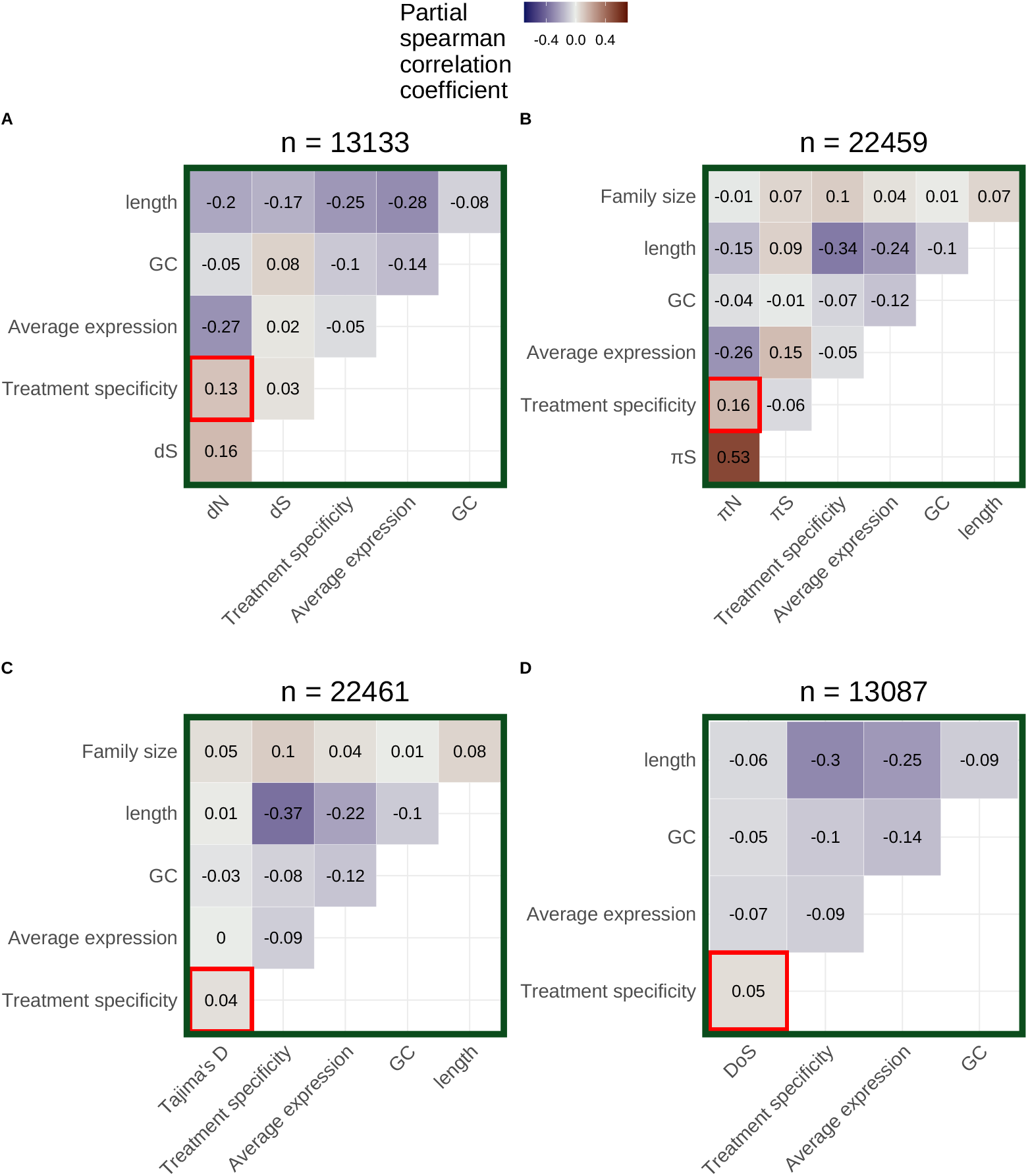
Partial correlations for (**A**) *dN*, (**B**) *π*_*N*_, (**C**) Tajima’s D, and (**D**) direction of selection (DoS) based on leaf tissue data subset after applying SVA. Data was further subset to include only treatment groups with data from more than one study before applying SVA. Average expression calculation excludes values ¡ 5 TPM. The number of genes included in each partial correlation analysis (n) is listed at the top of each heatmap.

## 4 Discussion

Our main finding is that genes with more treatment-specific expression patterns are, on average, under weaker selective constraint in *A. thaliana*. This is evident by treatment-specific genes generally having higher values of *π*_*N*_ and *dN*, but not higher values of Tajima’s D and DoS, compared to genes with more constitutive expression (Figures 4,5). Our result does not refute the possibility of strong positive selection on treatment-specific genes, as is the case for nucleotide binding site leucine rich repeat proteins (NBS-LRRs) in *A. thaliana* [51]. Rather, treatment-specific genes are simply under weaker selection on average compared to less treatment-specific genes. Altogether, this pattern is consistent with the hypothesis that a trade-off between the strength of selection and the treatment specificity of expression helps maintain variation in plasticity for *A. thaliana* [71, 79].

There are a few ways to think about the biological relevance of the correlations of treatment specificity with *π*_*N*_ and *dN*. First, the magnitude of treatment specificity’s correlation with *π*_*N*_ and *dN* was generally half the magnitude of average expression’s correlation with *π*_*N*_ and *dN* and similar to tissue specificity’s correlation with *π*_*N*_ and *dN*. Both tissue specificity and average expression are thought to be important determinants of protein evolution [7, 87], suggesting the comparable effects of treatment specificity may be important too. Second, the effect of treatment specificity on *π*_*N*_ and *dN* persisted even after simultaneously controlling for expression level, tissue specificity, gene length, GC content, and batch effects. Finally, the top 25% most treatment-specific genes in our dataset have average *dN* and *π*_*N*_ values nearly 2.5 times greater than the 25% least treatment-specific genes (*dN* = 0.025 vs 0.061; *π*_*N*_ = 0.0014 vs 0.0032), but relatively similar Tajima’s D and DoS values (Tajima’s D = -0.44 vs -0.43; DoS = -0.19 vs -0.14). These observations together suggest that treatment specificity is an important determinant of protein evolution.

This study disentangles several processes that were often difficult to resolve in previous research. First, many previous studies focus mainly on explaining trends in *dN/dS* [70, 27, *7*], but both relaxed negative selection and increased positive selection can lead to increases in *dN/dS*. To tease apart these two processes, we additionally investigated treatment specificity’s relationship with Tajima’s D and DoS. Treatment specificity’s weaker correlation with Tajima’s D and DoS, compared to *dN* and *π*_*N*_, suggests that relaxed negative selection plays a larger role than increased positive selection in explaining the high evolutionary rates of treatment-specific genes. Furthermore, measures of expression specificity are often highly correlated with expression level [70, 3, 30]. When calculating a gene’s expression level, we only included samples where said gene was expressed (TPM ¿ 5) to get an estimate of expression level that was still correlated with *dN* and *π*_*N*_, but was independent of expression specificity, allowing us to better disentangle these factors. Finally, previous studies have struggled to partition the factors that influence selection on genes in the presence of predictor variables with considerable error, such as expression level [21, 65, 90]. Error in expression measurements can often be attributed to unmeasured differences between RNA-sequencing experiments [44] and we accounted for these differences using SVA [45]. Even after SVA, treatment specificity was strongly correlated with *dN* and *π*_*N*_ (Figures 5A-B), suggesting our results are not an artifact of errors in expression measurement or combining expression data across many studies.

Surprisingly, nearly all genes in *A. thaliana* have some degree of treatment specificity in their expression (Figures 2A, S3), reflecting results of previous studies on tissue specificity [23]. The high prevalence of treatment specificity in our dataset is partly explained by batch effects because SVA significantly lowered the apparent treatment specificity of most genes (Figures S16B-S21B) and reduced within-treatment differentiation in PCA space (for example, see Figures S16C and S16D). This reduction in treatment-specificity likely happened because batch effects can include unrecorded between-treatment differences (e.g. the humidity of the growth chamber, light intensity, watering schedule, etc.). Controlling for these unrecorded between-treatment differences thus causes the expression of genes to be less treatment-specific. However, even after batch correction most genes still showed some degree of treatment specificity (Figures S16B-S21B), suggesting it is rare for a gene to be expressed at the same level across many environments.

We also observed that genes with higher treatment specificity generally belonged to larger gene families. We expected gene family size to correlate with selection because singleton and duplicated genes often evolve at different rates [33, 19]. Theory also suggests that gene duplication leads to relaxation of selection on duplicates, allowing for neo- and sub-functionalization [48, 1]. We could not investigate how gene family size correlates with *dN* or DoS because measuring these quantities requires identifying substitutions between orthologous genes. Thus, *dN* and *DoS* can only be reliably measured for 1:1 orthologs between *A. thaliana* and *A. lyrata*. However, *π*_*N*_ and Tajima’s D can be calculated for genes in larger families and we did observe persistent correlations between family size and Tajima’s D (For Figure 5C: *ρ* = 0.05, p-value = 3.1 *×* 10^−12^; also see Figures S6C-S15C, S28C-S33C). Altogether, these correlations suggest that processes of gene duplication, neofunctionalization, and subfunctionalization could be connected to evolving some degree of treatment specificity.

Gene length was generally the second most correlated factor with *dN* and *π*_*N*_ in our study, just behind average expression. This is consistent with previous work suggesting that longer proteins require more energy to synthesize and are thus under stronger selective constraints [76, 8, 23, 77]. However, while some previous studies in *A. thaliana* observe this same trend [7], others do not [70]. This discrepancy could be due to differences in how gene length is defined between studies. In this study, each gene’s length included coding sequence as well as introns and untranslated regions, whereas other studies break down gene length into individual features [7]. The goal of this study was not to understand differences in evolution between different gene features, so we included all gene features in our estimate of gene length. However, introns and untranslated regions experience different evolutionary patterns than coding sequences; for example, highly expressed genes being under selection for shorter introns [8, 23]. Therefore, future studies must clearly define even seemingly simple features like gene length to ensure that results are comparable across studies.

Although we focused on testing the idea that treatment specificity is responsible for relaxed negative selection in some genes, it is also possible that relaxed selection caused the evolution of treatment specificity. There is some evidence that relaxation of selection occurs before the evolution of expression specificity [32] and may better explain cases of neo- and subfunctionalization [48, 1]. Future experiments that look at the evolution of treatment specificity and sequence evolution across a broader phylogenetic scale may be helpful for determining the order of these processes.

In summary, this study investigates a trade-off between the treatment-specific expression of a gene and the strength of selection said gene experiences, which is hypothesized to limit plasticity evolution. Consistent with this hypothesis, genes in *A. thaliana* with more treatment-specific expression are under weaker selection compared to more evenly expressed genes. While we find that this trade-off exists, we could not dissect the direction of causality in the trade-off or determine how much this trade-off constrains plasticity evolution relative to other processes. However, these are exciting areas of future research. Future studies should ideally generate fully balanced datasets on gene expression acquired across natural environmental gradients. Taking these steps will contribute to a comprehensive understanding of the constraints on plasticity and protein evolution.

## 5 Data availability

All code for our bioinformatic workflows, data analysis, and figure creation can be found here:

https://github.com/milesroberts-123/arabidopsis-conditional-expression. The tissue type and treatment annotations for RNA-seq runs in our study can be found in Table S5. Genomic references as well as a table of expression specificity, nucleotide diversity, and substitution rate values estimated for all *A. thaliana* genes included in this manuscript’s analyses is available at: XXX (url inserted at publication). The genome assembly and annotation used in this study was originally downloaded from Phytozome: https://phytozome-next.jgi.doe.gov/.

## 6 Acknowledgments

We would like to thank the following individuals for helpful comments on the initial drafts of this manuscript: Robert VanBuren, Shinhan Shiu, Stephen I. Wright, Nathan Catlin, Rebecca Panko, Asia Hightower, Maya Wilson Brown, Sophie Buysse, and Mia Stevens. We would like to further thank the two anonymous peer-reviewers whose comments significantly improved later drafts of this manuscript. Finally, we want to acknowledge the hundreds of scientists throughout the world that measured gene expression in *A. thaliana* and made their data publicly available. Without them, this work would not be possible.

