## Supplemental Figures for "Weaker selection on genes with treatment-specific expression consistent with a limit on plasticity evolution in *Arabidopsis thaliana*"

#### Contents

**1 General data curation** **2**

**2 Partial correlations on overall dataset** **6**

**3 Partial correlations on data subset by tissue type after SVA** **11**

**4 PCA before and after SVA on data subset by tissue type** **16**

**5 Legends - PCA before and after SVA on data subset by tissue type** **22**

**6 Partial correlations, balanced subset after SVA** **26**

**7 Effect of omitting low-expression values on treatment-specificity vs expression level correlation** **32**

**8 Simulating level-specificity correlation** **37**

### 1 General data curation

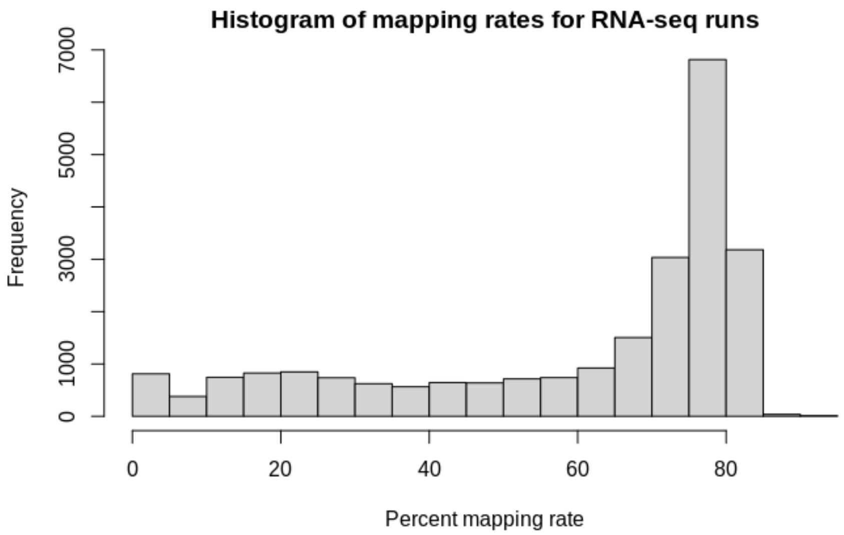


Figure S1: Histogram of mapping rates for RNA-seq runs

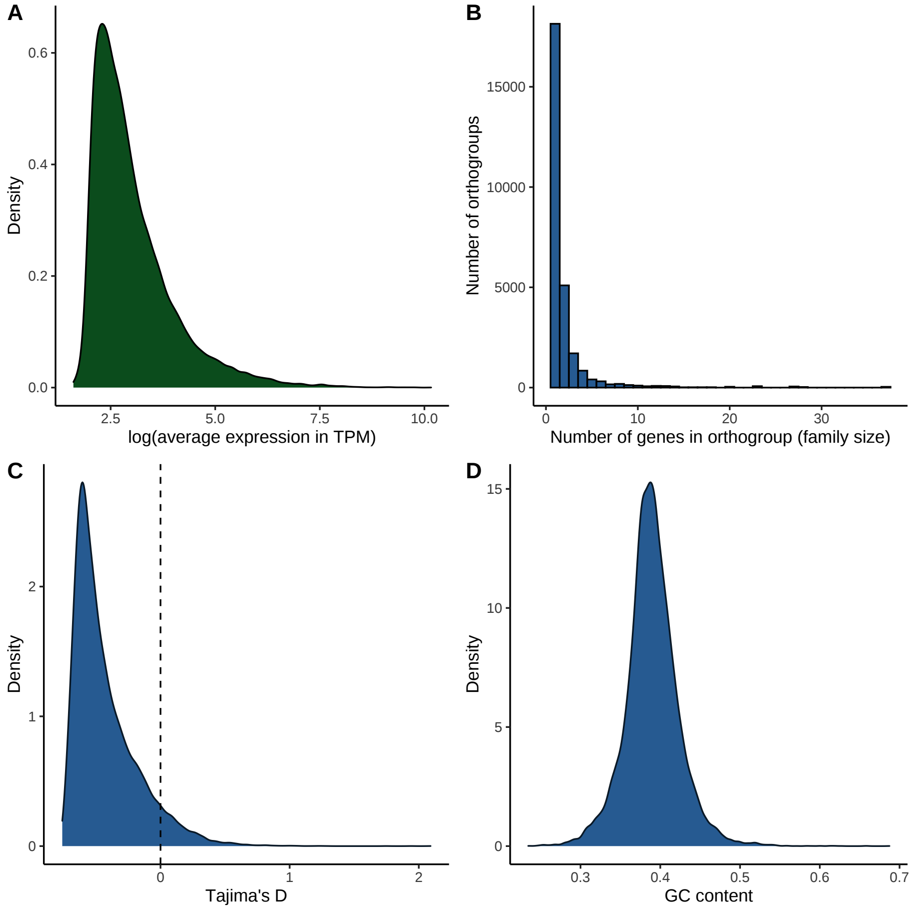


Figure S2: More histograms of other key variables for the genes included in this study.

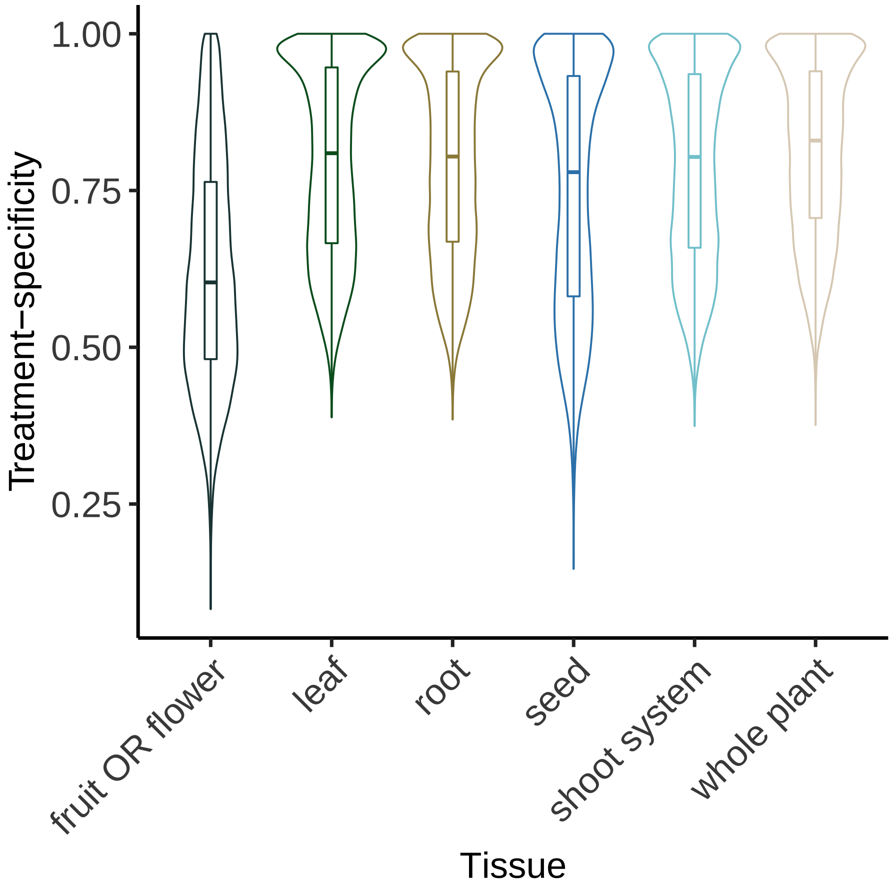


Figure S3: Violin plot of treatment-specificity in gene expression for six tissue categories.

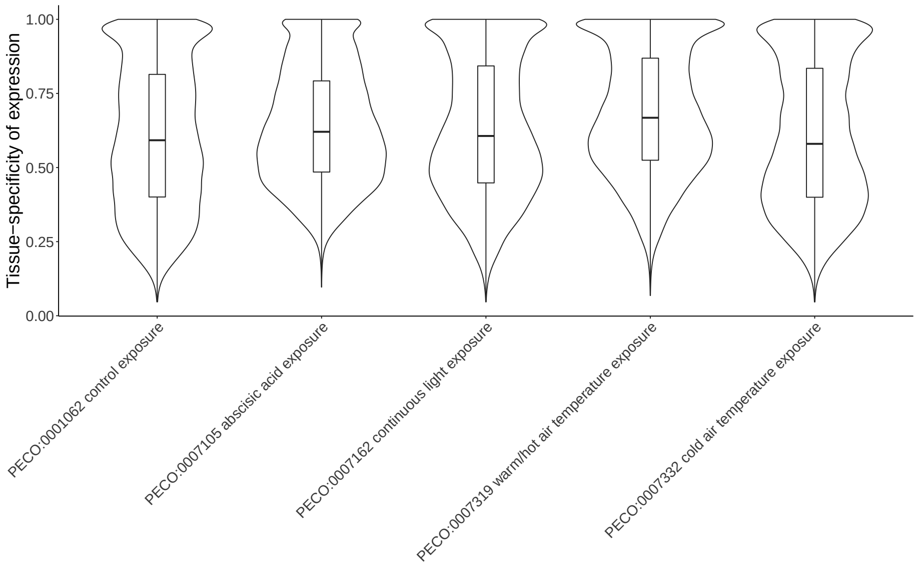


Figure S4: Violin of tissue-specificity in gene expression by treatment type. Only the five treatments shown had samples from all six tissue types.

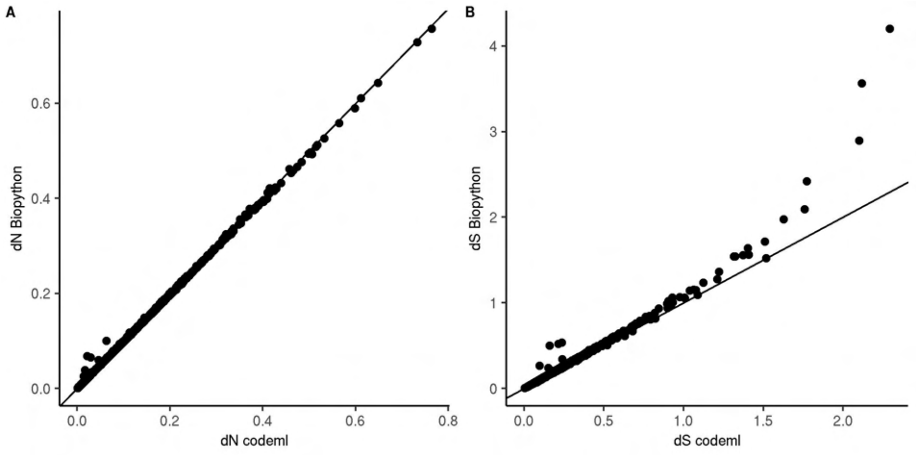


Figure S5: Comparing $dN$ and $dS$ values returned from codeml and the custom biopython script used in this study. Line shows exact match between codeml and biopython methods. Genes with saturating divergence ($dS > 1$) were excluded from partial correlation analyses.

### 2 Partial correlations on overall dataset

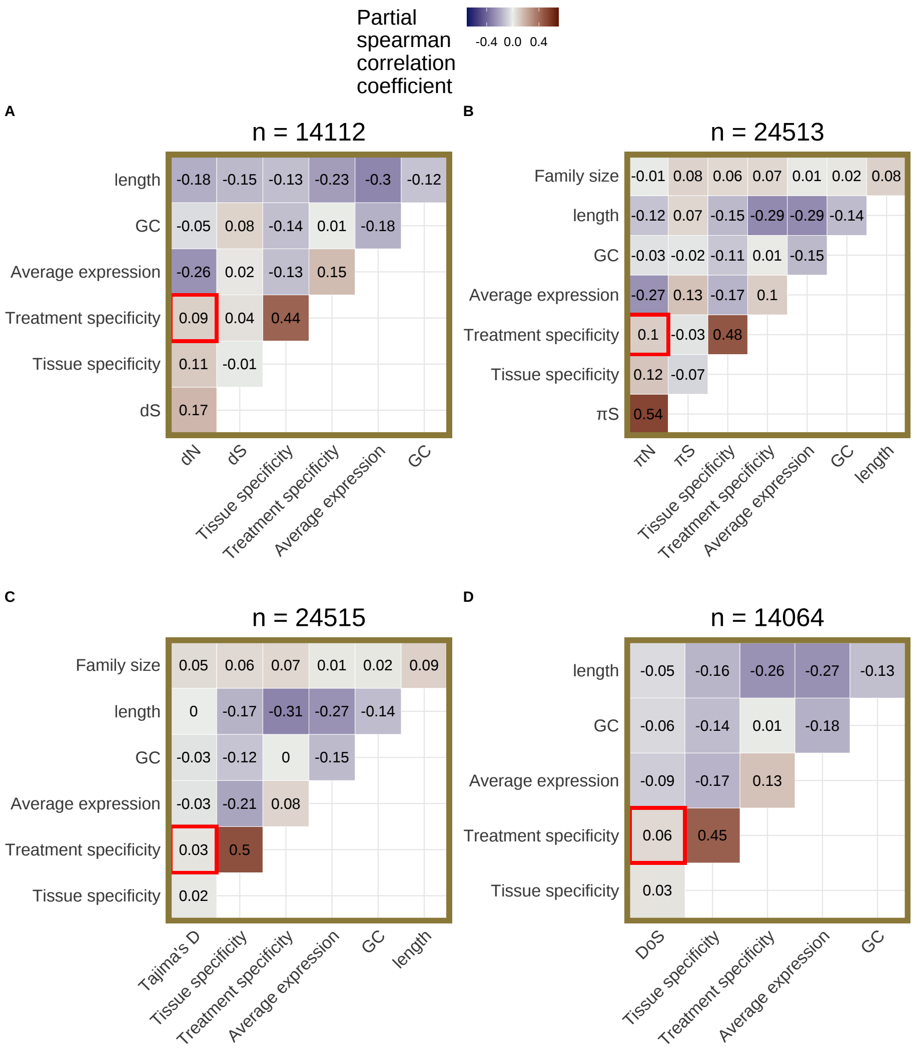


Figure S6: Partial correlations for (**A**) $dN$, (**B**) $\pi_N$, (**C**) Tajima's D, and (**D**) direction of selection (DoS) based on root data. Average expression excludes values $< 5$ TPM and was calculated using only root tissue samples. Tissue-specificity was calculated using only control runs across all tissue categories. Treatment-specificity was calculated using only root tissue runs. The number of genes included in each partial correlation analysis (n) is listed at the top of each heatmap.

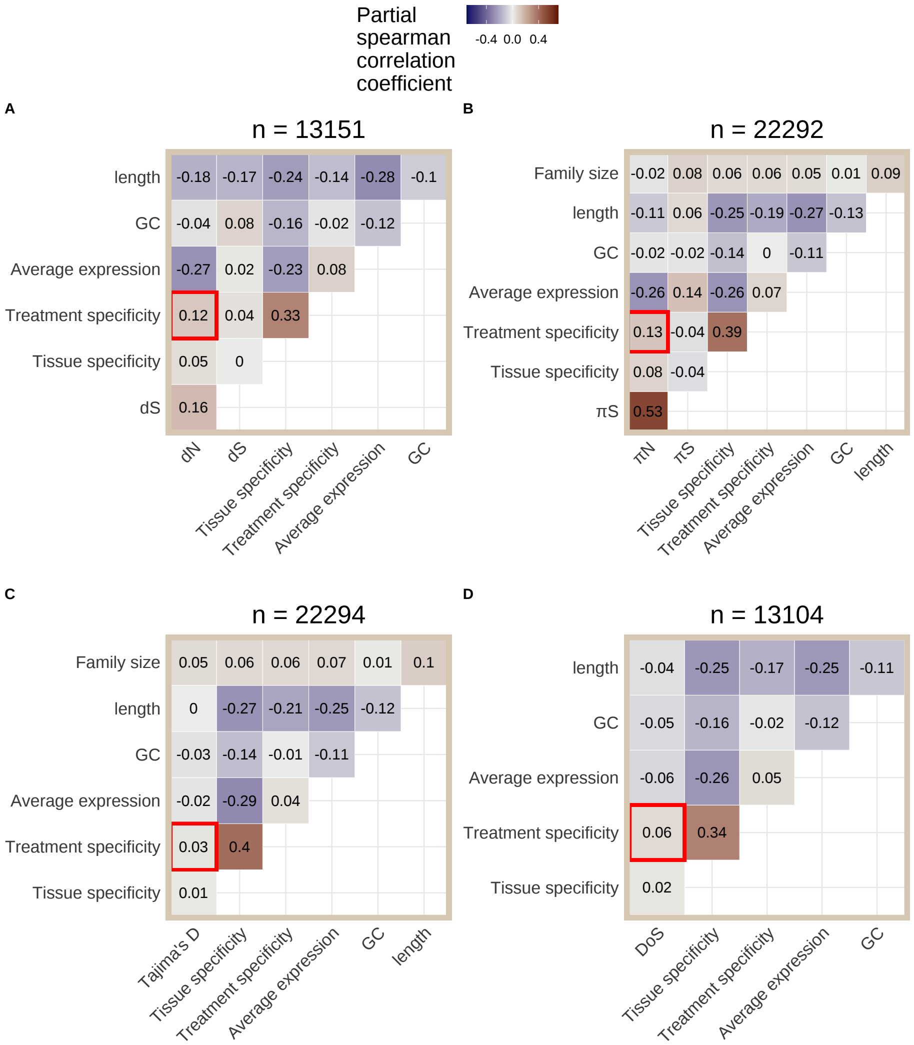


Figure S7: Partial correlations for (**A**) $dN$, (**B**) $\pi_N$, (**C**) Tajima's D, and (**D**) direction of selection (DoS) based on whole plant data. Average expression excludes values $< 5$ TPM and was calculated using only whole plant tissue samples. Tissue-specificity was calculated using only control runs across all tissue categories. Treatment-specificity was calculated using only whole plant tissue runs. The number of genes included in each partial correlation analysis (n) is listed at the top of each heatmap.

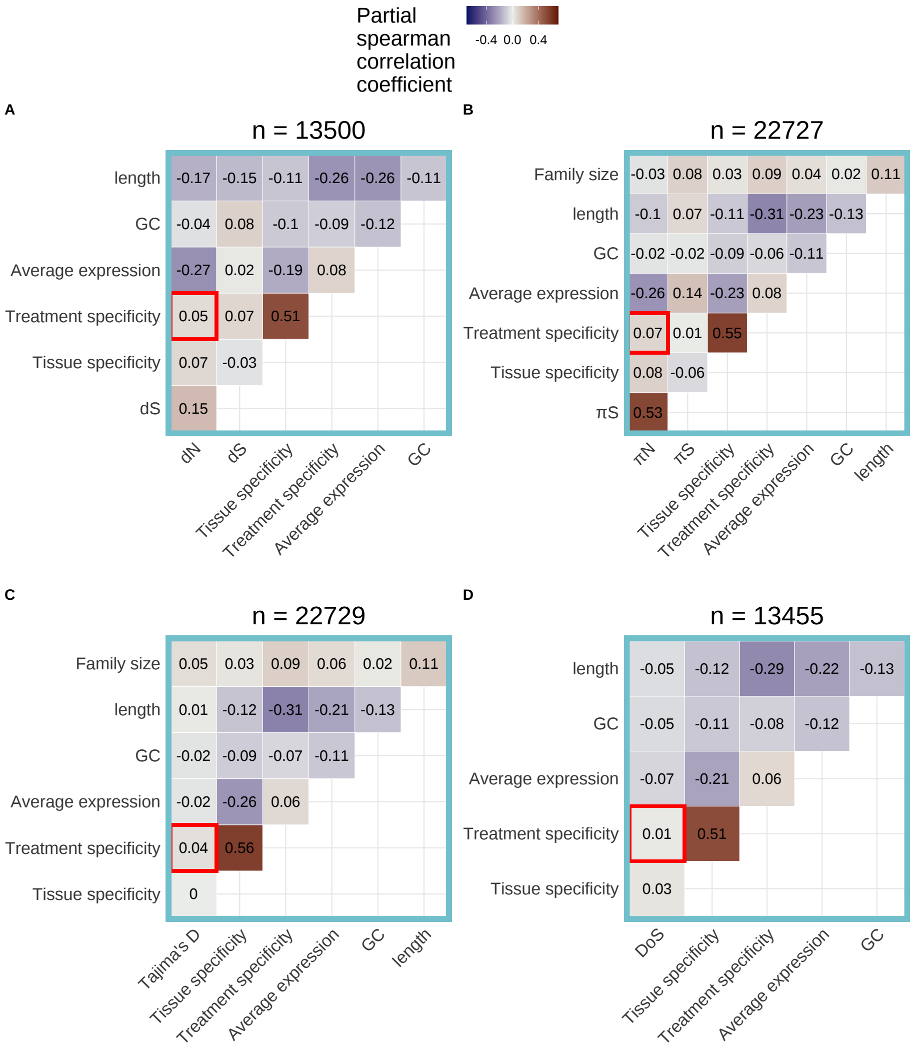


Figure S8: Partial correlations for (**A**) $dN$, (**B**) $\pi_N$, (**C**) Tajima's D, and (**D**) direction of selection (DoS) based on shoot data. Average expression excludes values $< 5$ TPM and was calculated using only shoot tissue samples. Tissue-specificity was calculated using only control runs across all tissue categories. Treatment-specificity was calculated using only shoot tissue runs. The number of genes included in each partial correlation analysis (n) is listed at the top of each heatmap.

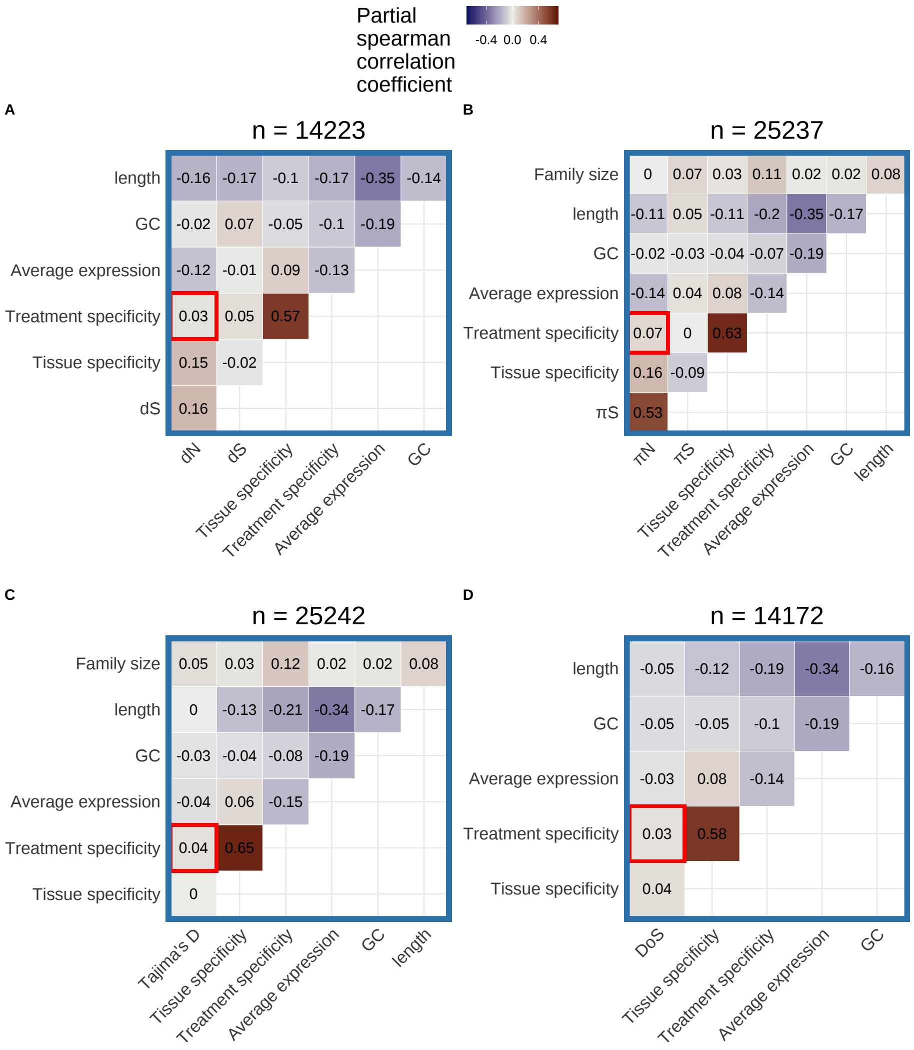


Figure S9: Partial correlations for (**A**) $dN$, (**B**) $\pi_N$, (**C**) Tajima's D, and (**D**) direction of selection (DoS) based on seed data. Average expression excludes values $<$ 5 TPM and was calculated using only seed tissue samples. Tissue-specificity was calculated using only control runs across all tissue categories. Treatment-specificity was calculated using only seed tissue runs. The number of genes included in each partial correlation analysis (n) is listed at the top of each heatmap.

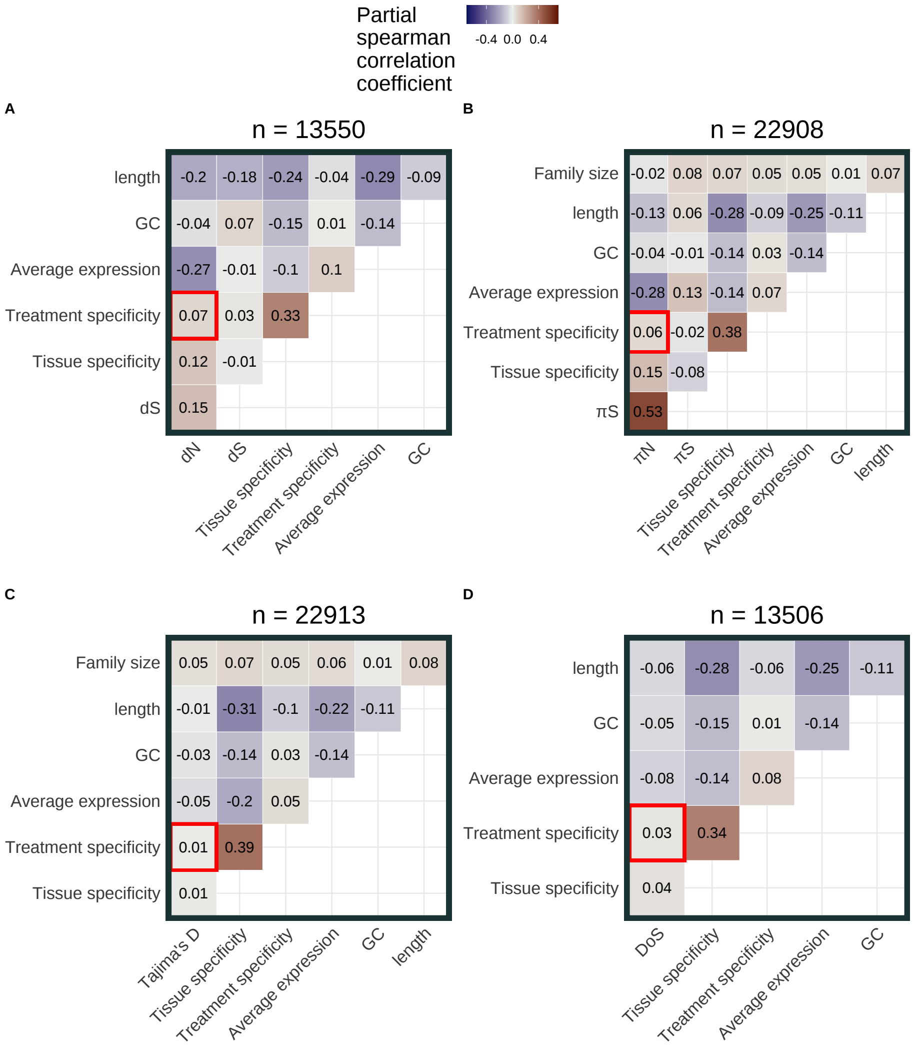

Figure S10: Partial correlations for (**A**) $dN$, (**B**) $\pi_N$, (**C**) Tajima's D, and (**D**) direction of selection (DoS) based on flower and fruit data. Average expression excludes values < 5 TPM and was calculated using only fruit and flower tissue samples. Tissue-specificity was calculated using only control runs across all tissue categories. Treatment-specificity was calculated using only fruit and flower tissue runs. The number of genes included in each partial correlation analysis (n) is listed at the top of each heatmap.

### 3 Partial correlations on data subset by tissue type after SVA

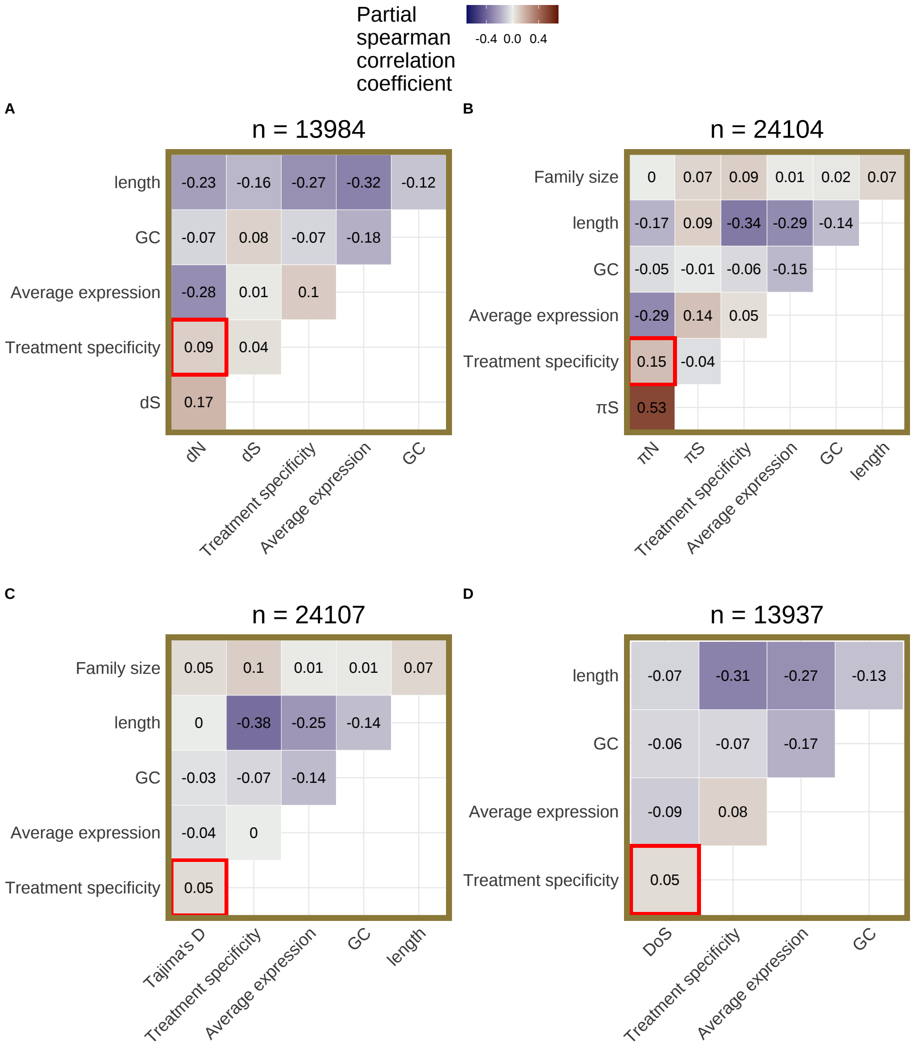


Figure S11: Partial correlations for (**A**) $dN$, (**B**) $\pi_N$, (**C**) Tajima's D, and (**D**) direction of selection (DoS) based on root tissue data after applying SVA. Data was further subset to include only treatment groups with data from more than one study before applying SVA. The number of genes included in each partial correlation analysis (n) is listed at the top of each heatmap.

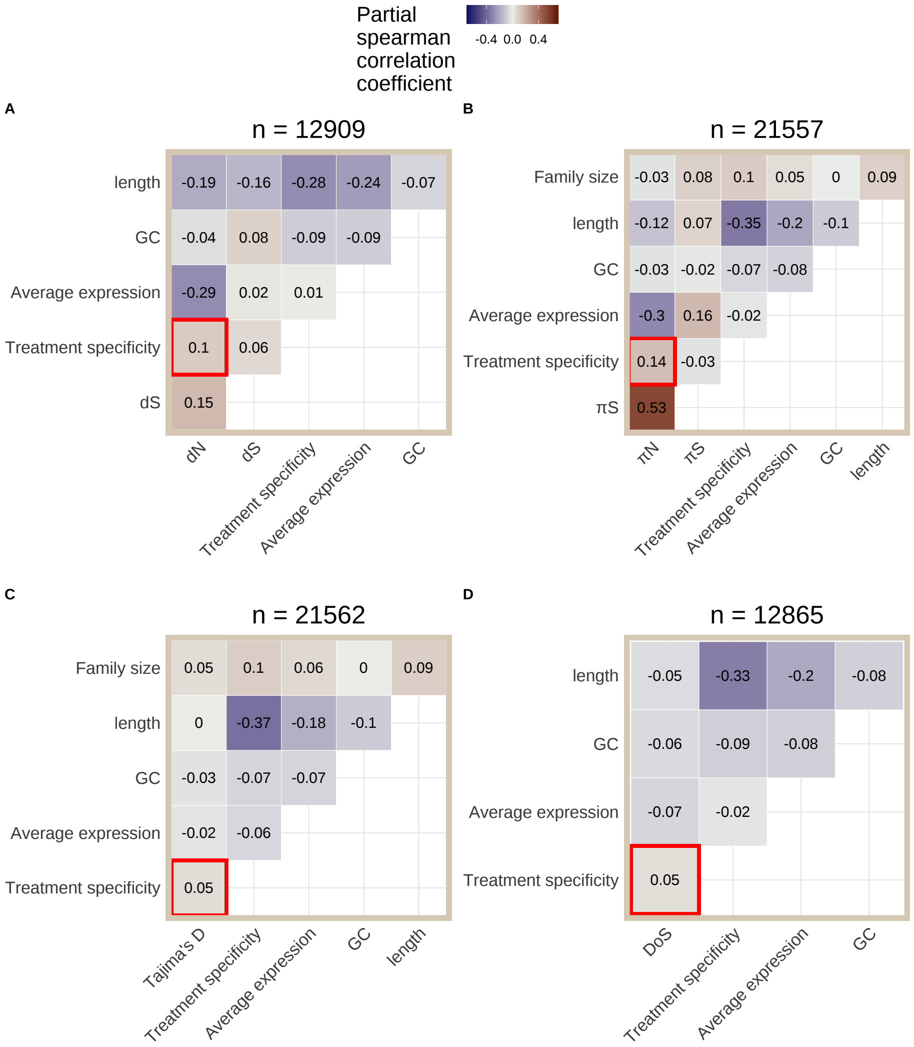


Figure S12: Partial correlations for (**A**) $dN$, (**B**) $\pi_N$, (**C**) Tajima's D, and (**D**) direction of selection (DoS) based on whole plant tissue data after applying SVA. Data was further subset to include only treatment groups with data from more than one study before applying SVA. The number of genes included in each partial correlation analysis (n) is listed at the top of each heatmap.

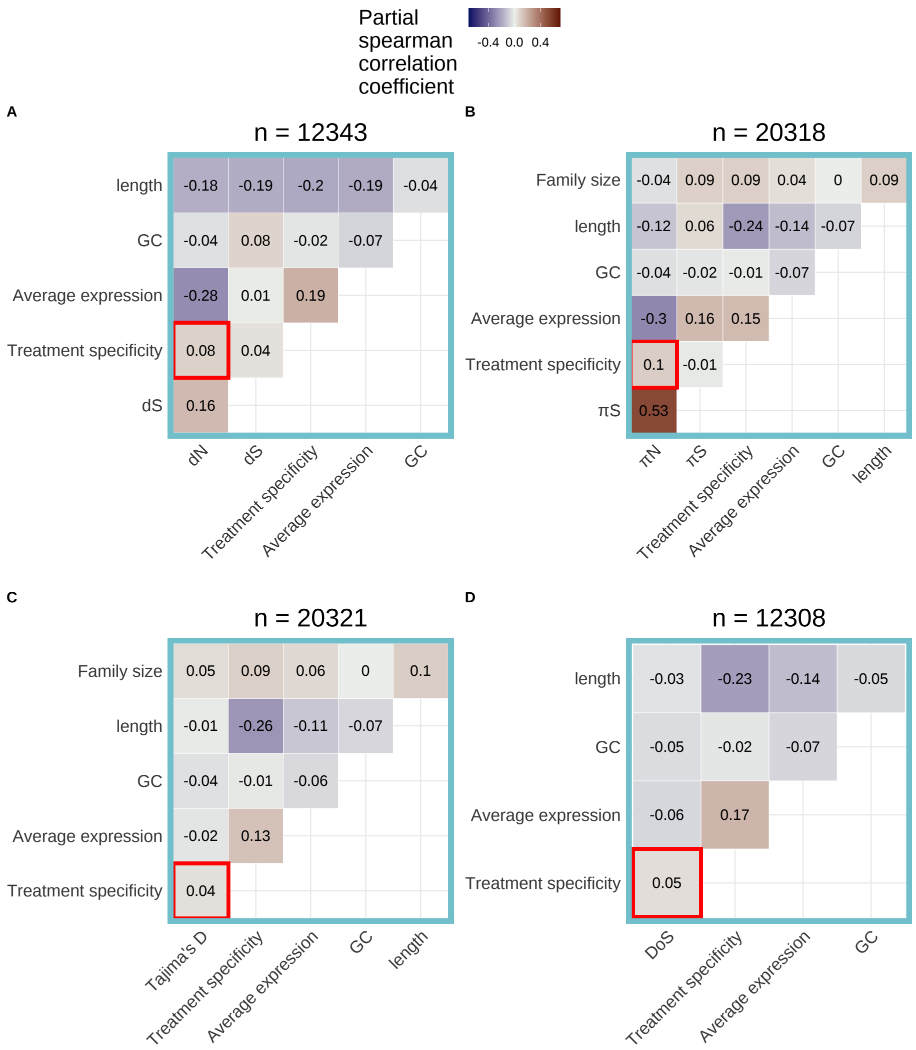


Figure S13: Partial correlations for (**A**) $dN$, (**B**) $\pi_N$, (**C**) Tajima's D, and (**D**) direction of selection (DoS) based on shoot tissue data after applying SVA. Data was further subset to include only treatment groups with data from more than one study before applying SVA. The number of genes included in each partial correlation analysis (n) is listed at the top of each heatmap.

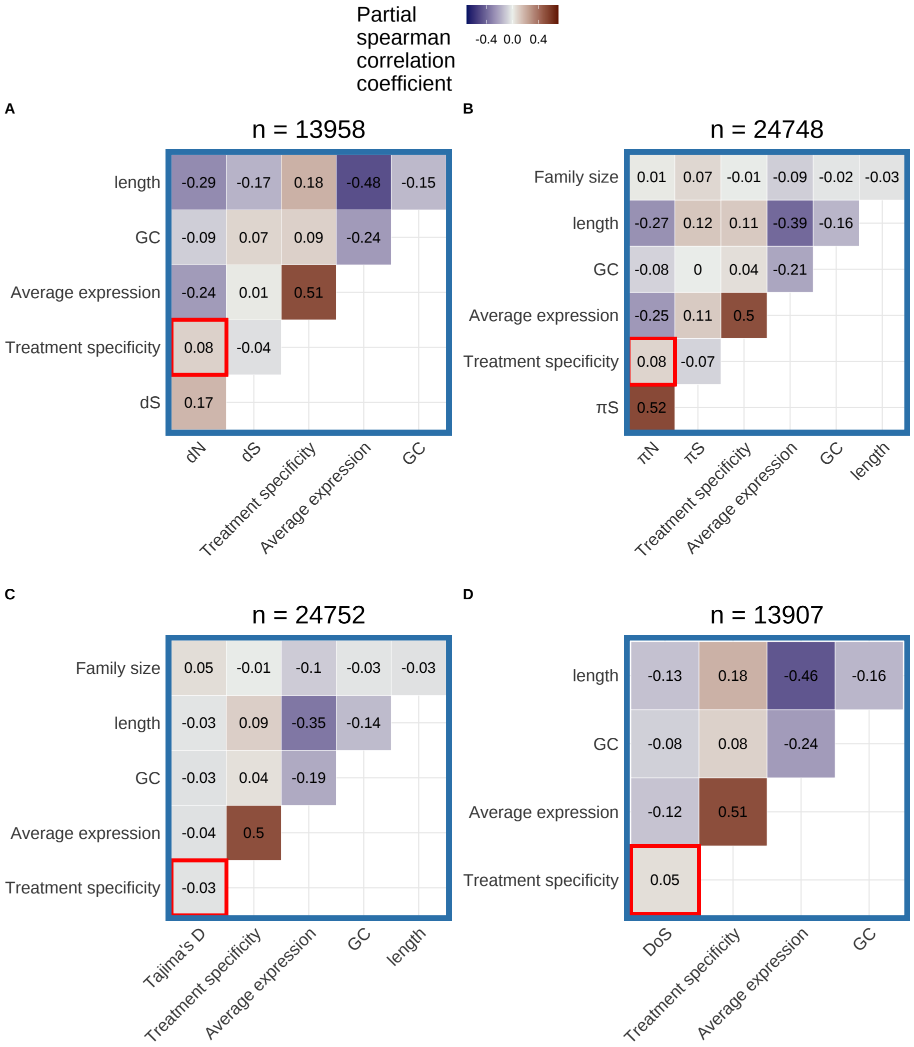


Figure S14: Partial correlations for (**A**) $dN$, (**B**) $\pi_N$, (**C**) Tajima's D, and (**D**) direction of selection (DoS) based on seed tissue data after applying SVA. Data was further subset to include only treatment groups with data from more than one study before applying SVA. The number of genes included in each partial correlation analysis (n) is listed at the top of each heatmap.

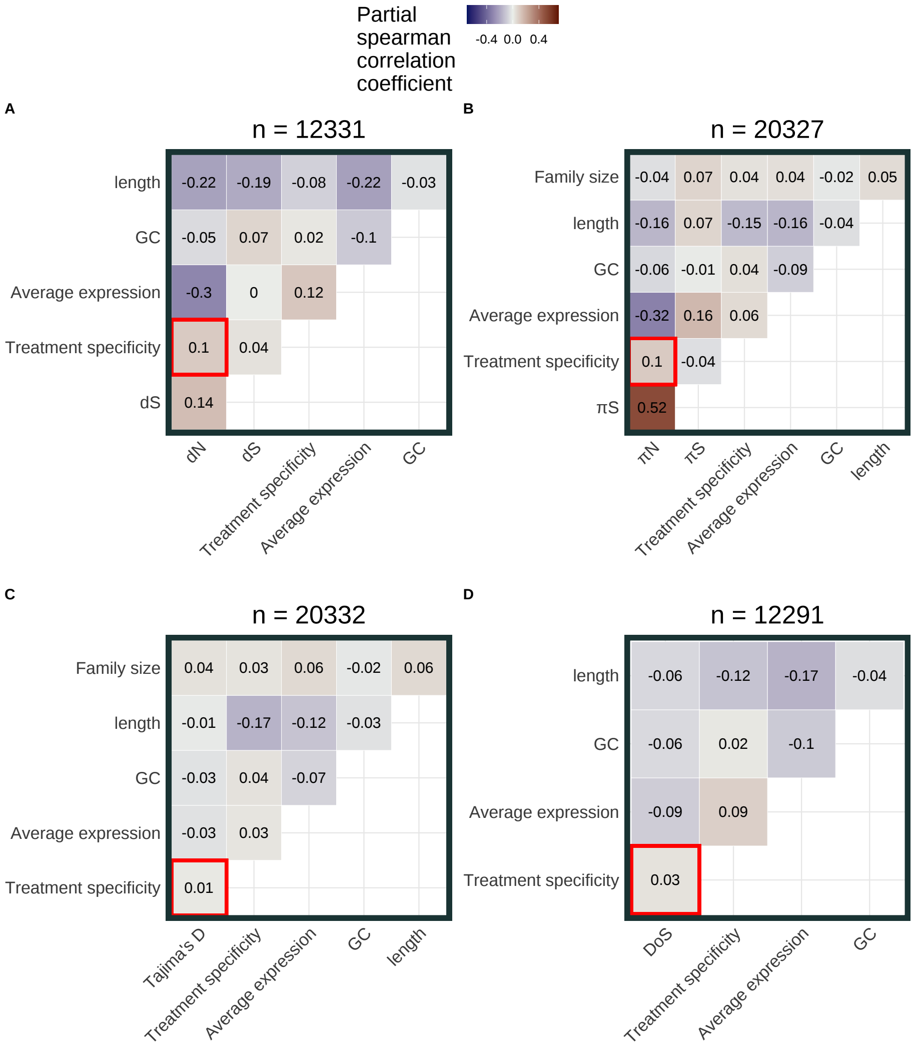


Figure S15: Partial correlations for (**A**) $dN$, (**B**) $\pi_N$, (**C**) Tajima's D, and (**D**) direction of selection (DoS) based on flower and fruit tissue data after applying SVA. Data was further subset to include only treatment groups with data from more than one study before applying SVA. The number of genes included in each partial correlation analysis (n) is listed at the top of each heatmap.

#### 4 PCA before and after SVA on data subset by tissue type

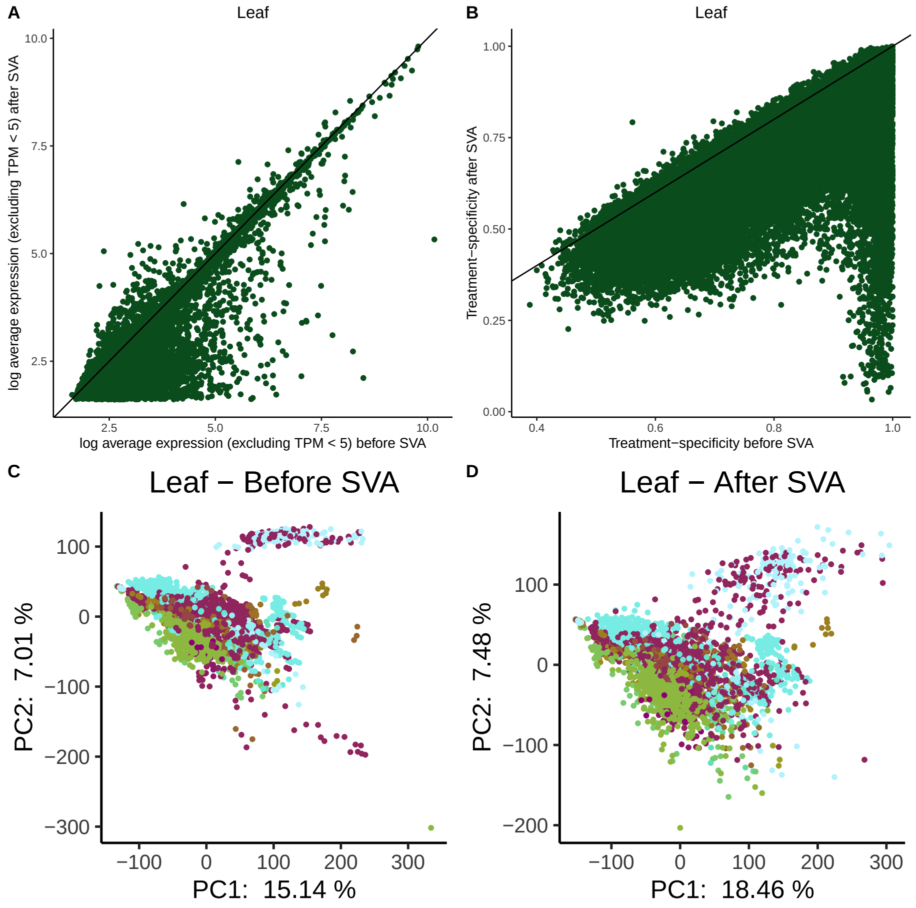


Figure S16: Effect of surrogate variable analysis on structure of leaf tissue data. (**A**) Effect of SVA on average expression. (**B**) Effect of SVA on treatment-specificity. (**C**) Principal component analysis of leaf tissue data before SVA. (**D**) Principal component analysis of leaf tissue data after SVA. Color in the PCA plots represent different experimental treatments - legend is Figure S22.

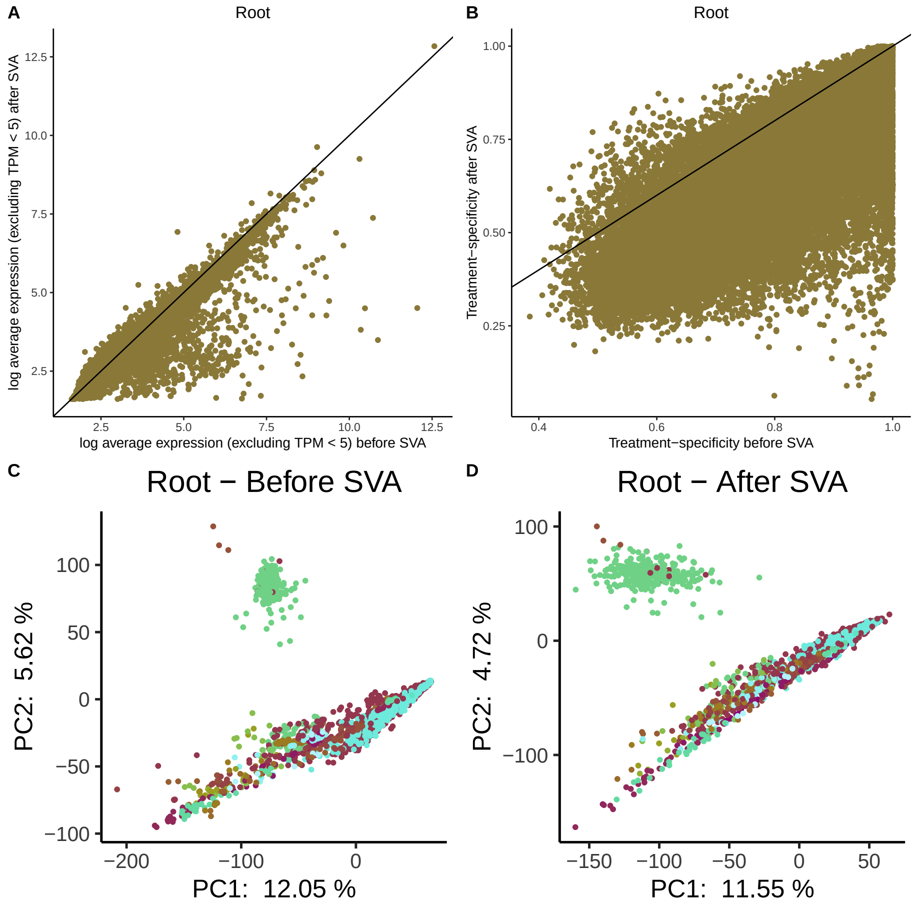


Figure S17: Effect of surrogate variable analysis on structure of root tissue data. (**A**) Effect of SVA on average expression. (**B**) Effect of SVA on treatment-specificity. (**C**) Principal component analysis of root tissue data before SVA. (**D**) Principal component analysis of root tissue data after SVA.Color in the PCA plots represent different experimental treatments - legend is Figure S23.

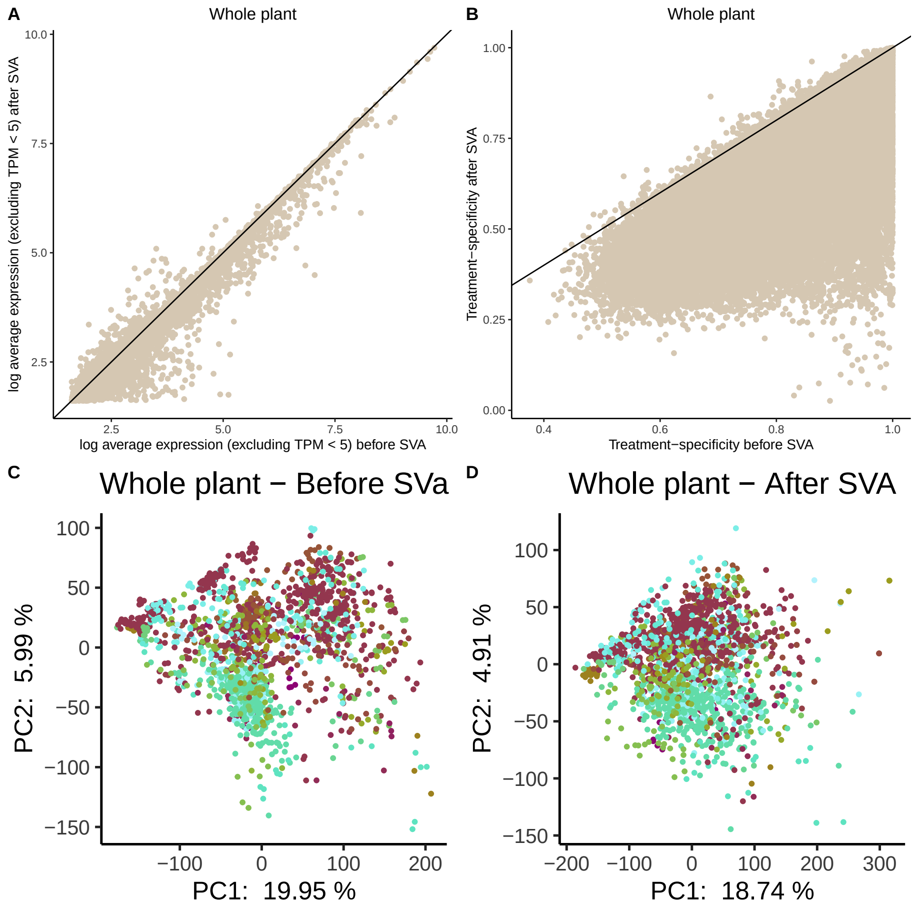


Figure S18: Effect of surrogate variable analysis on structure of whole plant tissue data. (**A**) Effect of SVA on average expression. (**B**) Effect of SVA on treatment-specificity. (**C**) Principal component analysis of whole plant tissue data before SVA. (**D**) Principal component analysis of whole plant tissue data after SVA.Color in the PCA plots represent different experimental treatments - legend is Figure S24.

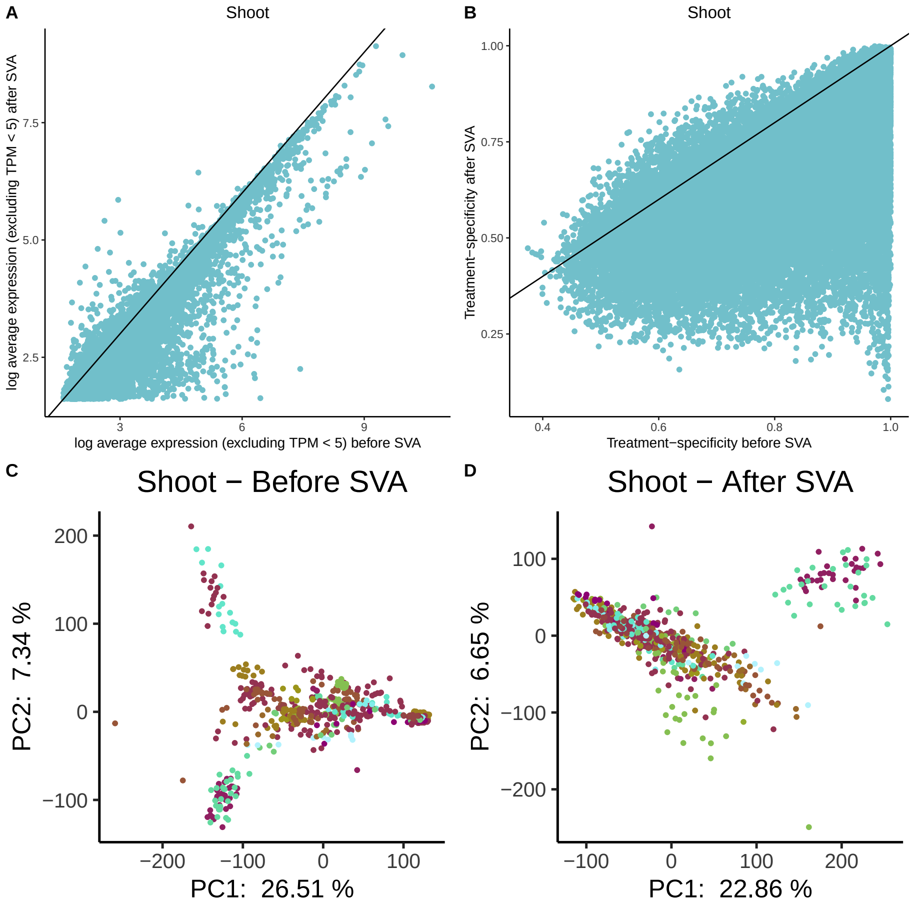


Figure S19: Effect of surrogate variable analysis on structure of shoot tissue data. (**A**) Effect of SVA on average expression. (**B**) Effect of SVA on treatment-specificity. (**C**) Principal component analysis of shoot tissue data before SVA. (**D**) Principal component analysis of shoot tissue data after SVA. Color in the PCA plots represent different experimental treatments - legend is Figure S25.

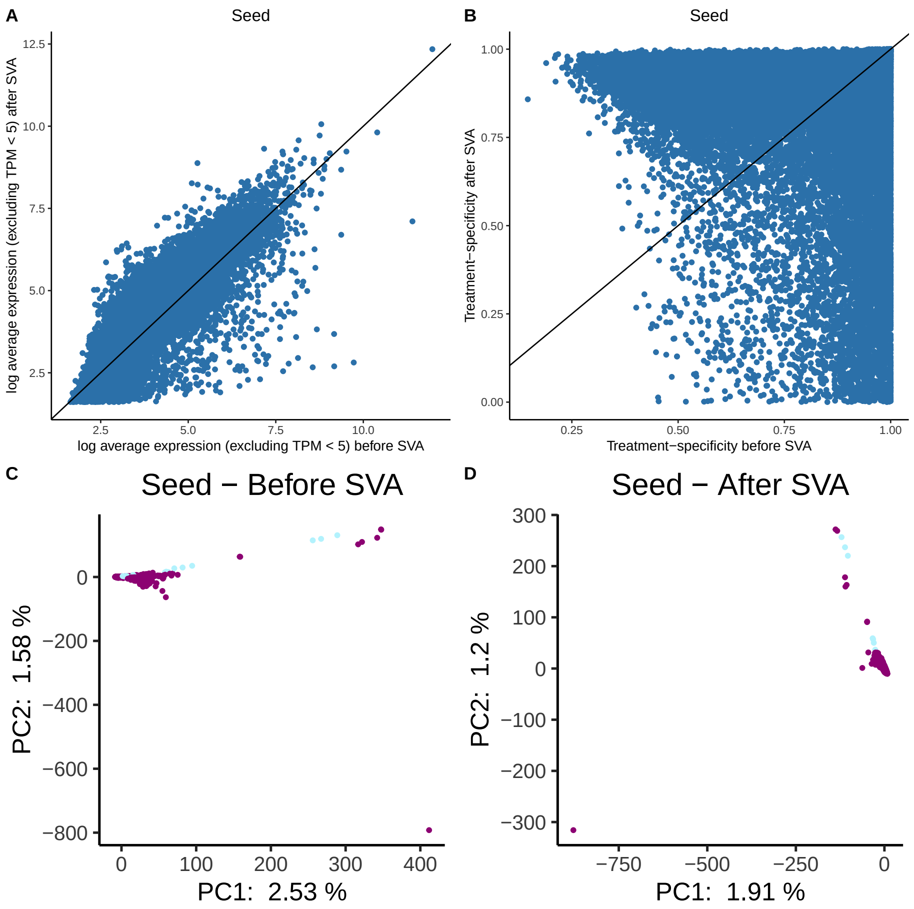


Figure S20: Effect of surrogate variable analysis on structure of seed tissue data. (**A**) Effect of SVA on average expression. (**B**) Effect of SVA on treatment-specificity. (**C**) Principal component analysis of seed tissue data before SVA. (**D**) Principal component analysis of seed tissue data after SVA. Color in the PCA plots represent different experimental treatments - legend is Figure S26.

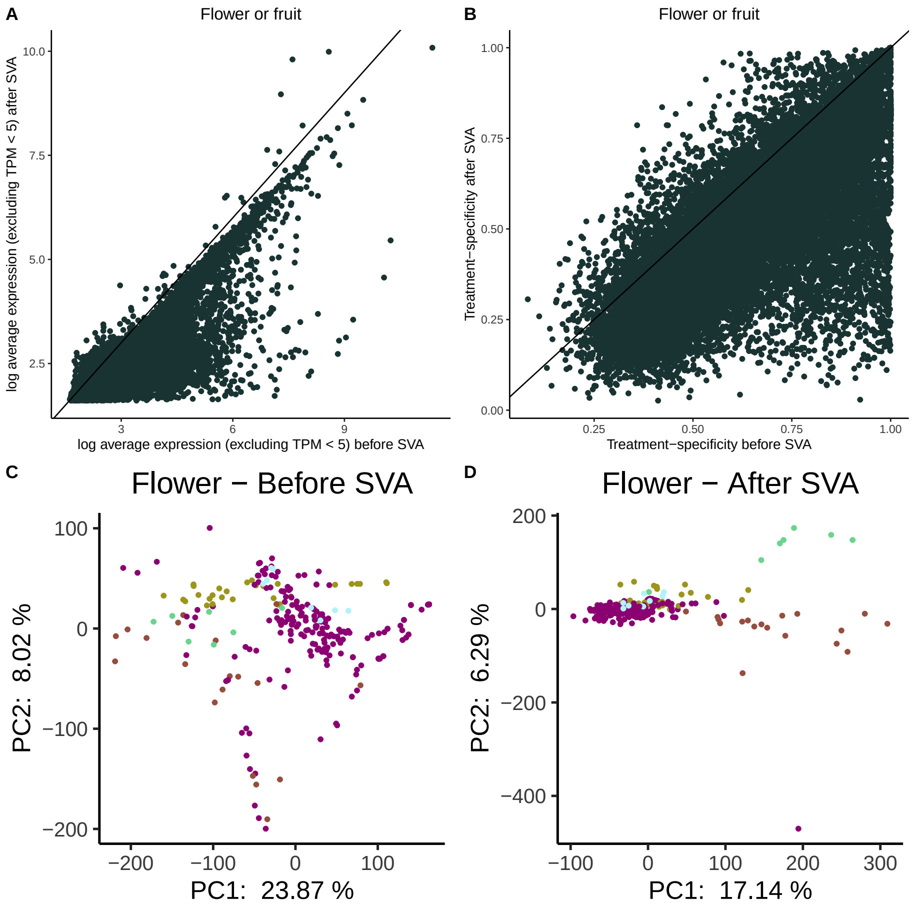


Figure S21: Effect of surrogate variable analysis on structure of fruit and flower tissue data. (**A**) Effect of SVA on average expression. (**B**) Effect of SVA on treatment-specificity. (**C**) Principal component analysis of fruit and flower tissue data before SVA. (**D**) Principal component analysis of fruit and flower tissue data after SVA. Color in the PCA plots represent different experimental treatments - legend is Figure S27.

### 5 Legends - PCA before and after SVA on data subset by tissue type

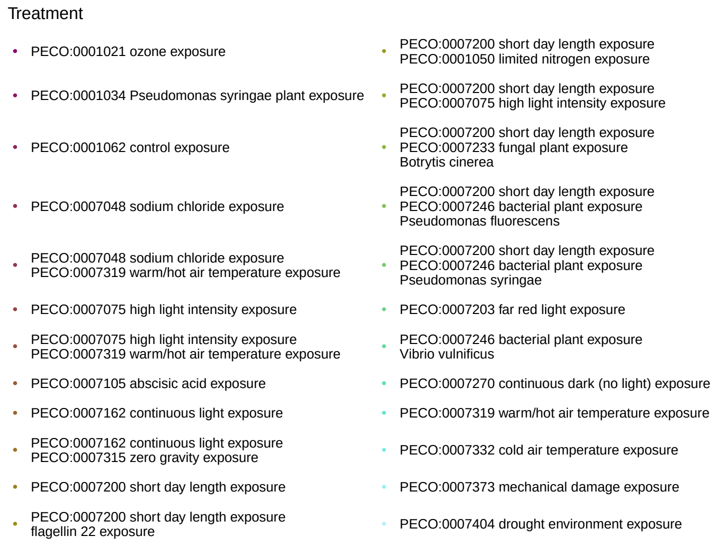


Figure S22: Legend for PCA before and after SVA, leaf data

- PECO:0001046 limited phosphate exposure
- PECO:0001050 limited nitrogen exposure
- PECO:0001062 control exposure
- PECO:0007162 continuous light exposure
- PECO:0007162 continuous light exposure<br>PECO:0007315 zero gravity exposure
- PECO:0007165 growth hormone exposure<br>ACC
- PECO:0007200 short day length exposure
- PECO:0007200 short day length exposure<br>flagellin 22 exposure
- PECO:0007200 short day length exposure<br>limited copper exposure
- PECO:0007200 short day length exposure<br>PECO:0001046 limited phosphate exposure
- PECO:0007200 short day length exposure<br>PECO:0001049 limited magnesium exposure
- PECO:0007200 short day length exposure<br>PECO:0007397 phosphorus nutrient exposure
- PECO:0007242 iron nutrient exposure
- PECO:0007246 bacterial plant exposure
- PECO:0007284 nitrogen macronutrient exposure
- PECO:0007319 warm/hot air temperature exposure
- PECO:0007373 mechanical damage exposure

Figure S23: Legend for PCA before and after SVA, root data

Treatment

- flagellin 22 exposure
- PECO:0001036 chemical stress exposure<br>tunicamycin
- PECO:0001038 osmotic stress exposure<br>mannitol
- PECO:0001046 limited phosphate exposure
- PECO:0001062 control exposure
- PECO:0007048 sodium chloride exposure
- PECO:0007105 abscisic acid exposure
- PECO:0007162 continuous light exposure
- PECO:0007162 continuous light exposure<br>PECO:0007001 UV−B light exposure
- PECO:0007162 continuous light exposure<br>PECO:0007048 sodium chloride exposure
- PECO:0007162 continuous light exposure<br>PECO:0007075 high light intensity exposure
- PECO:0007162 continuous light exposure<br>PECO:0007189 chemical exposure<br>DMSO
- PECO:0007162 continuous light exposure<br>PECO:0007319 warm/hot air temperature exposure
- PECO:0007189 chemical exposure<br>DMSO
- PECO:0007196 light exposure
- PECO:0007200 short day length exposure
- PECO:0007200 short day length exposure<br>flagellin 22 exposure
- PECO:0007200 short day length exposure<br>PECO:0007319 warm/hot air temperature exposure
- PECO:0007200 short day length exposure<br>PECO:0007332 cold air temperature exposure
- PECO:0007207 red light exposure
- PECO:0007246 bacterial plant exposure<br>PECO:0007397 phosphorus nutrient exposure
- PECO:0007270 continuous dark (no light) exposure
- PECO:0007319 warm/hot air temperature exposure
- PECO:0007332 cold air temperature exposure
- PECO:0007397 phosphorus nutrient exposure
- PECO:0007404 drought environment exposure

Figure S24: Legend for PCA before and after SVA, whole plant data

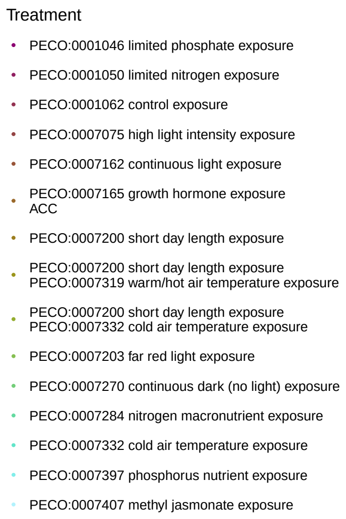


Figure S25: Legend for PCA before and after SVA, shoot data

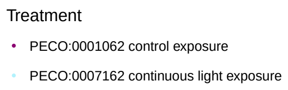


Figure S26: Legend for PCA before and after SVA, seed data

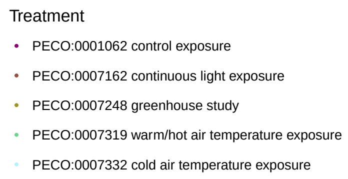


Figure S27: Legend for PCA before and after SVA, flower data

### 6 Partial correlations, balanced subset after SVA

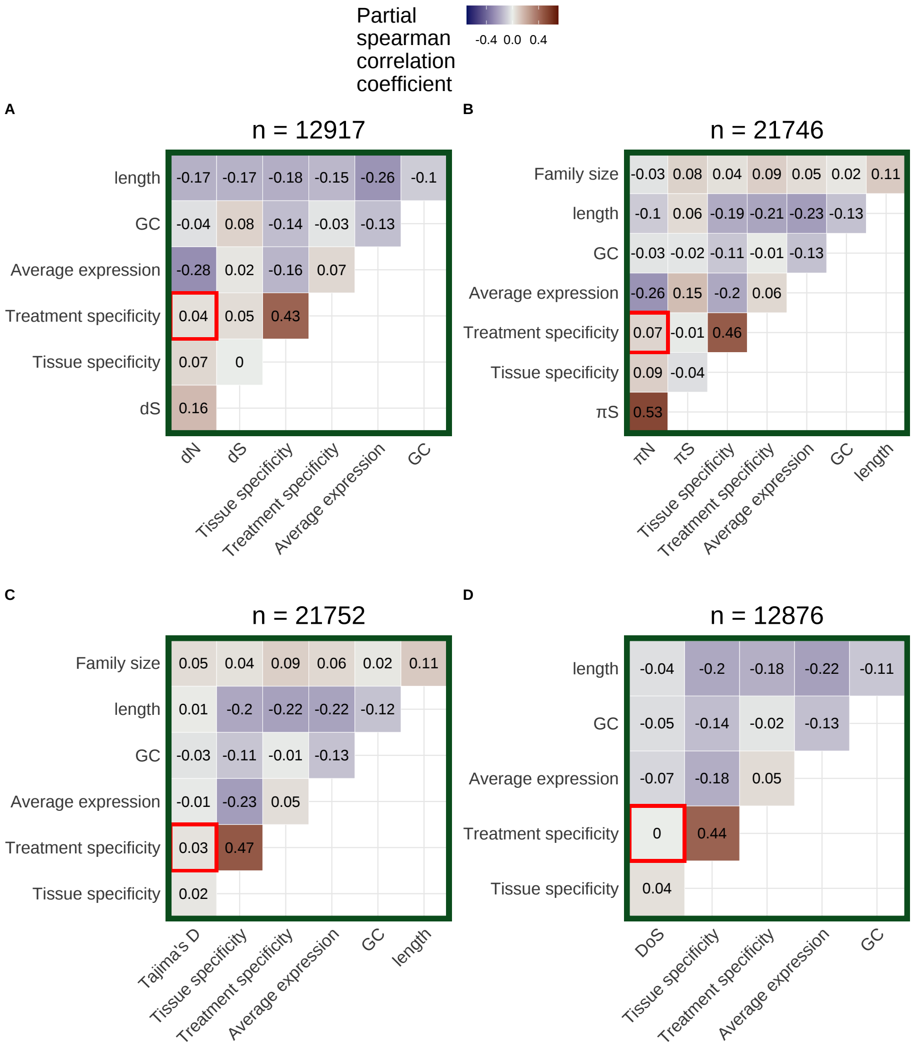


Figure S28: Partial correlations for (**A**) $dN$, (**B**) $\pi_N$, (**C**) Tajima's D, and (**D**) direction of selection (DoS) after applying SVA. Average expression and tissue specificity are calculated using only leaf tissue data. The number of genes included in each partial correlation analysis (n) is listed at the top of each heatmap.

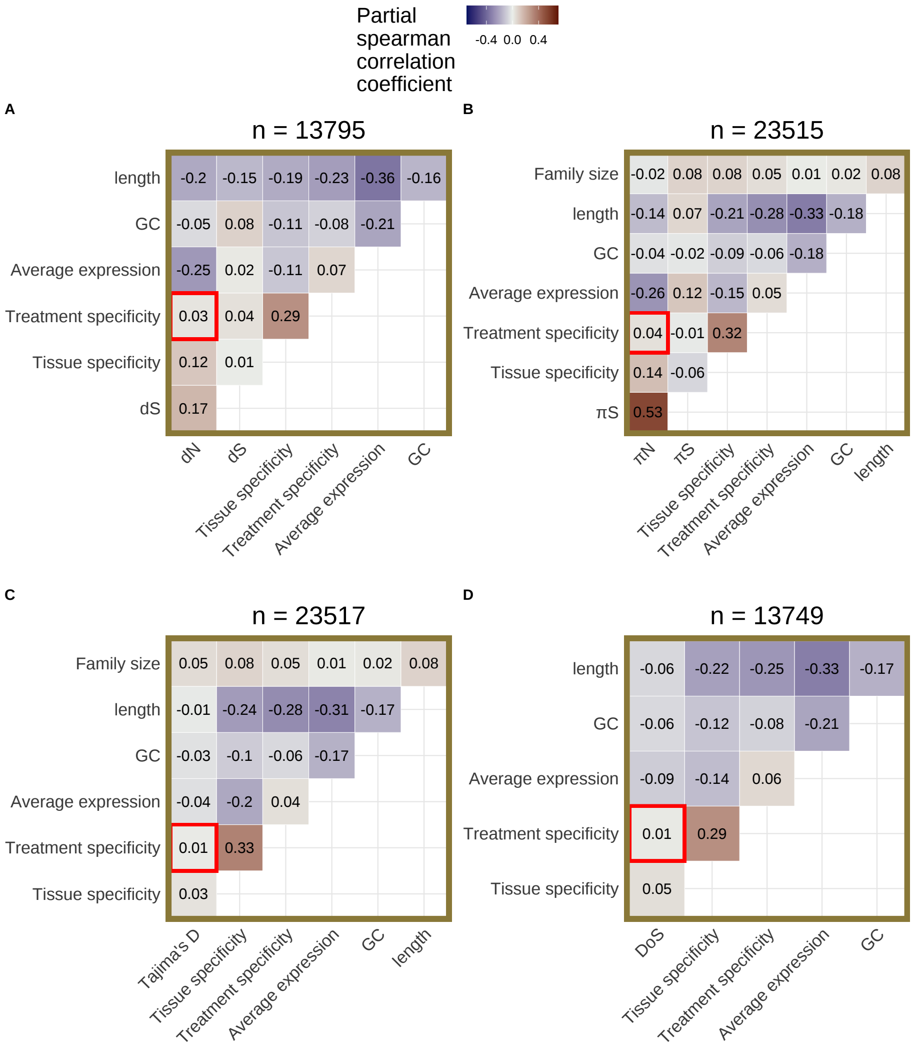


Figure S29: Partial correlations for (**A**) $dN$, (**B**) $\pi_N$, (**C**) Tajima's D, and (**D**) direction of selection (DoS) after applying SVA. Average expression and tissue specificity are calculated using only root tissue data. The number of genes included in each partial correlation analysis (n) is listed at the top of each heatmap.

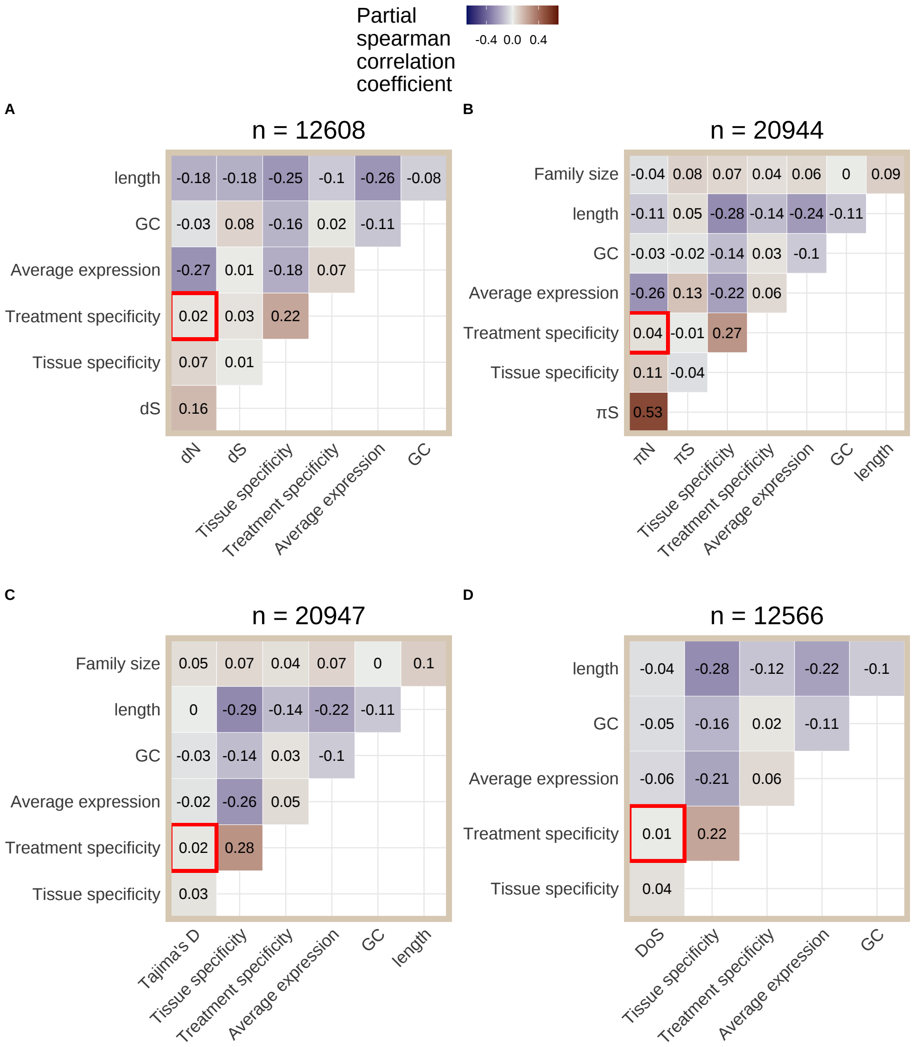


Figure S30: Partial correlations for (**A**) $dN$, (**B**) $\pi_N$, (**C**) Tajima's D, and (**D**) direction of selection (DoS) after applying SVA. Average expression and tissue specificity are calculated using only whole plant tissue data. The number of genes included in each partial correlation analysis (n) is listed at the top of each heatmap.

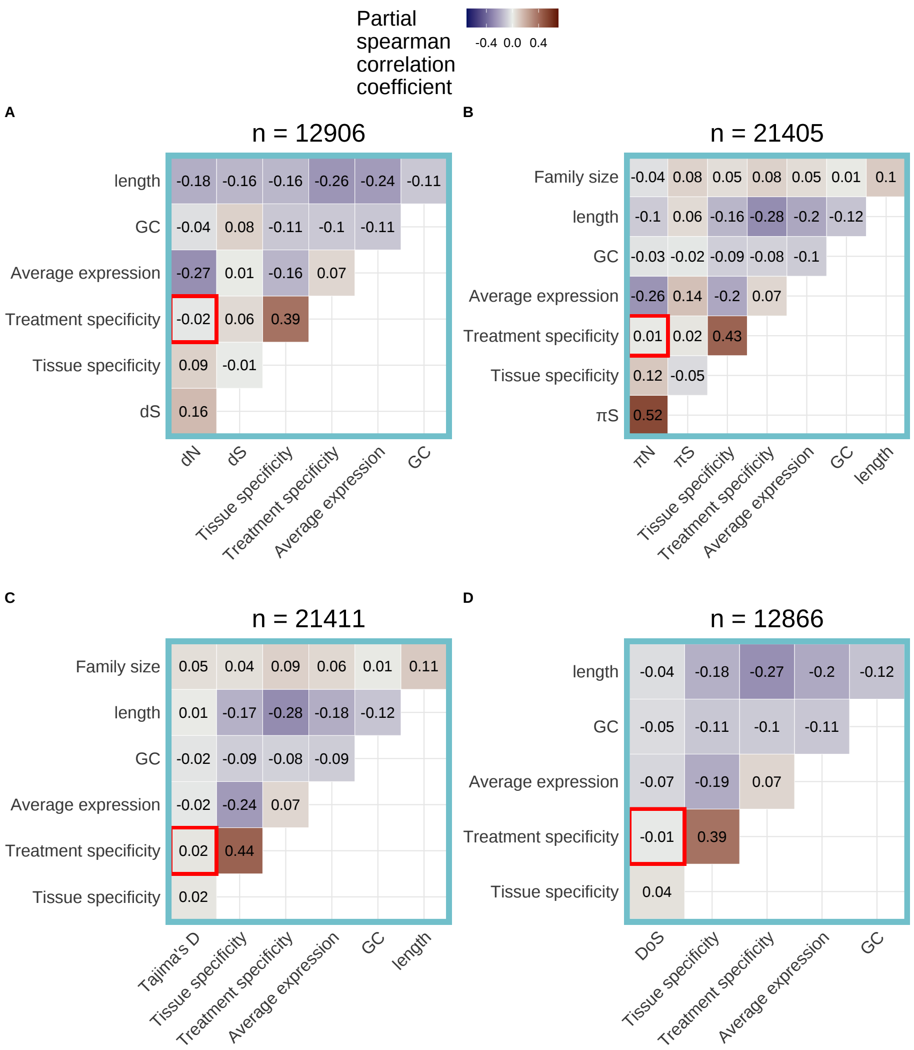


Figure S31: Partial correlations for (**A**) $dN$, (**B**) $\pi_N$, (**C**) Tajima's D, and (**D**) direction of selection (DoS) after applying SVA. Average expression and tissue specificity are calculated using only shoot tissue data. The number of genes included in each partial correlation analysis (n) is listed at the top of each heatmap.

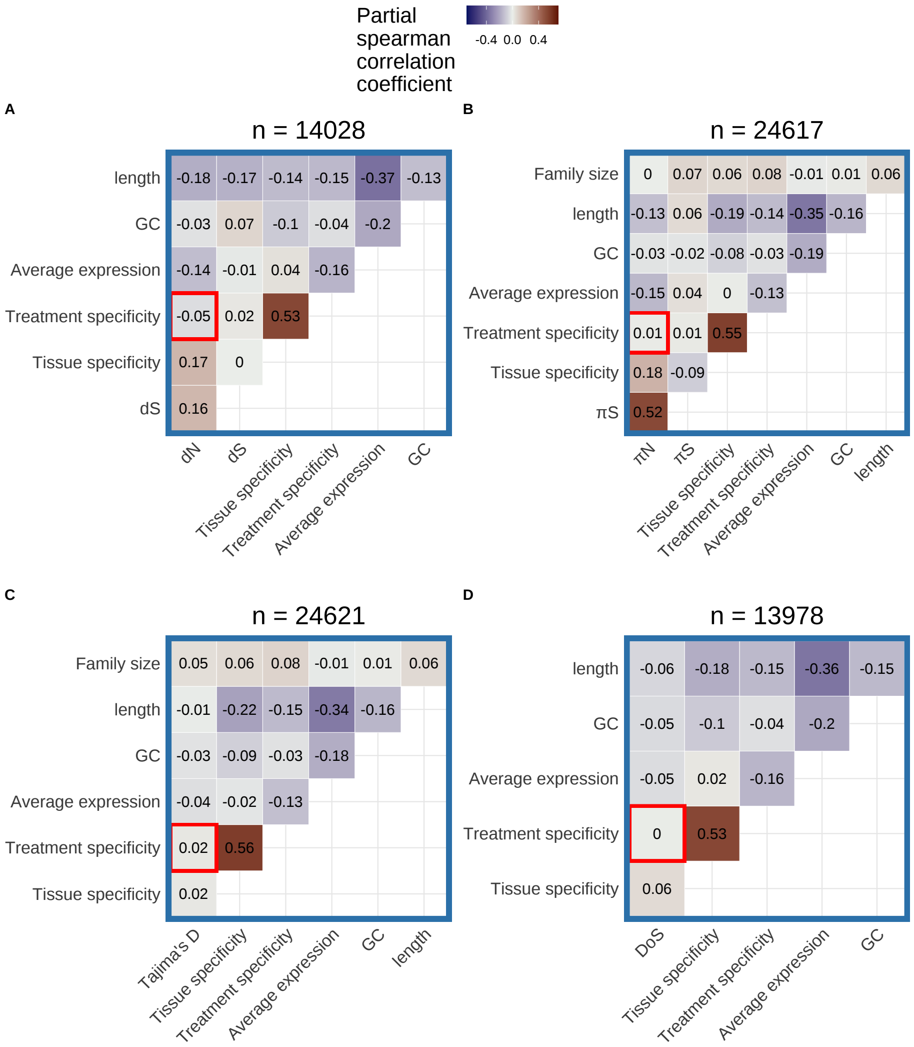


Figure S32: Partial correlations for (**A**) $dN$, (**B**) $\pi_N$, (**C**) Tajima's D, and (**D**) direction of selection (DoS) after applying SVA. Average expression and tissue specificity are calculated using only seed tissue data. The number of genes included in each partial correlation analysis (n) is listed at the top of each heatmap.

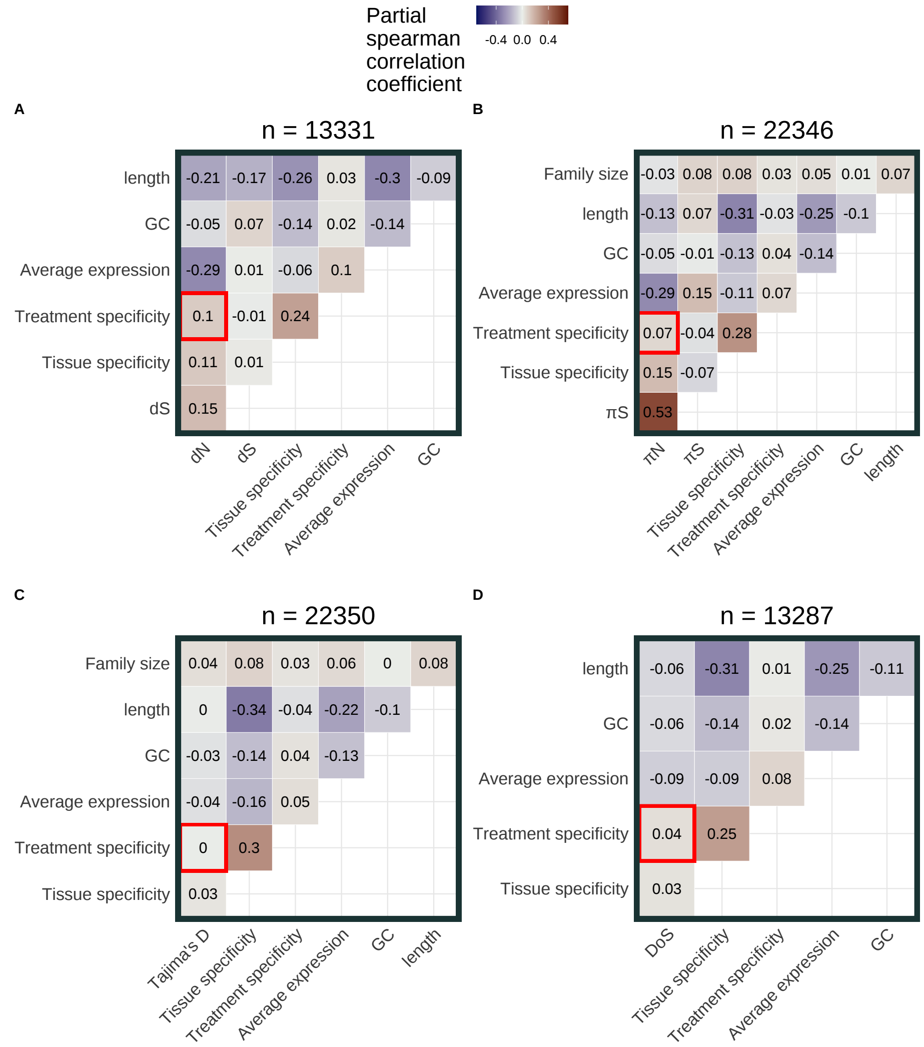


Figure S33: Partial correlations for (**A**) $dN$, (**B**) $\pi_N$, (**C**) Tajima's D, and (**D**) direction of selection (DoS) after applying SVA. Average expression and tissue specificity are calculated using only fruit and flower tissue data. The number of genes included in each partial correlation analysis (n) is listed at the top of each heatmap.

### 7 Effect of omitting low-expression values on treatment-specificity vs expression level correlation

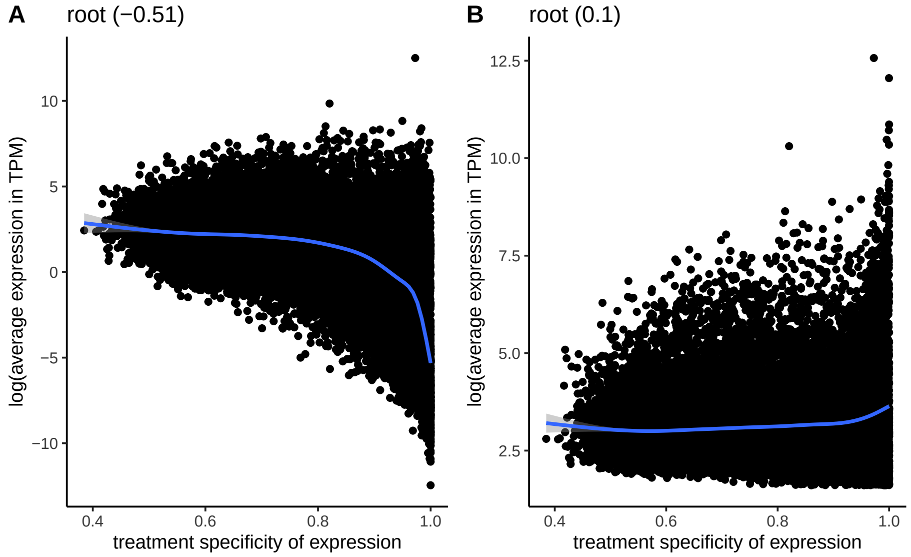


Figure S34: Correlation between average expression in transcripts per million (TPM) and treatment specificity of genes when low expression values (< 5 TPM) are included (**A**) vs excluded (**B**). Expression level and treatment specificity were calculated using only data from root tissue samples. Line is a smoothing line with 95 % confidence intervals and values in parentheses give spearman correlation.

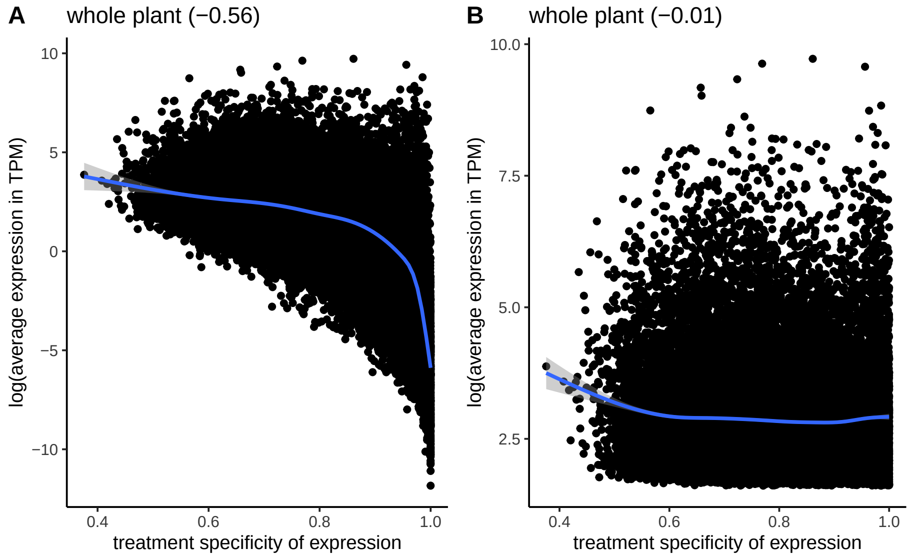


Figure S35: Correlation between average expression in transcripts per million (TPM) and treatment specificity of genes when low expression values (< 5 TPM) are included (**A**) vs excluded (**B**). Expression level and treatment specificity were calculated using only data from whole plant tissue samples. Line is a smoothing line with 95 % confidence intervals and values in parentheses give spearman correlation.

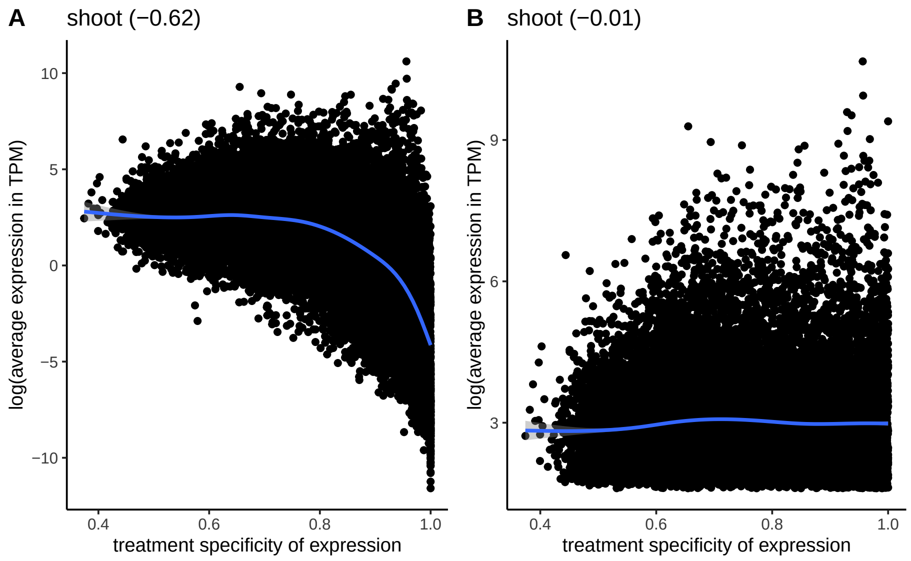


Figure S36: Correlation between average expression in transcripts per million (TPM) and treatment specificity of genes when low expression values (< 5 TPM) are included (**A**) vs excluded (**B**). Expression level and treatment specificity were calculated using only data from shoot tissue samples. Line is a smoothing line with 95 % confidence intervals and values in parentheses give spearman correlation.

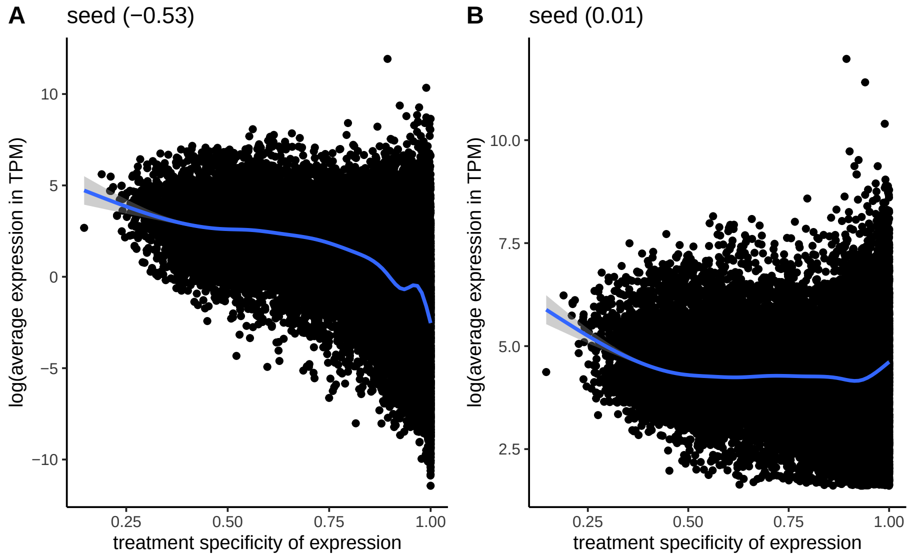


Figure S37: Correlation between average expression in transcripts per million (TPM) and treatment specificity of genes when low expression values (< 5 TPM) are included (**A**) vs excluded (**B**). Expression level and treatment specificity were calculated using only data from seed tissue samples. Line is a smoothing line with 95 % confidence intervals and values in parentheses give spearman correlation.

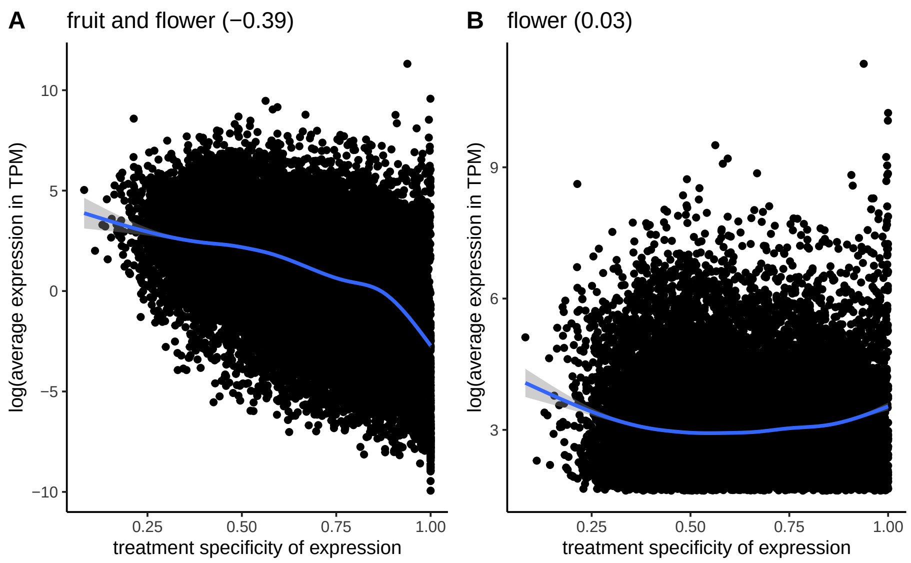


Figure S38: Correlation between average expression in transcripts per million (TPM) and treatment specificity of genes when low expression values (< 5 TPM) are included (**A**) vs excluded (**B**). Expression level and treatment specificity were calculated using only data from fruit and flower tissue samples. Line is a smoothing line with 95 % confidence intervals and values in parentheses give spearman correlation.

#### 8 Simulating level-specificity correlation

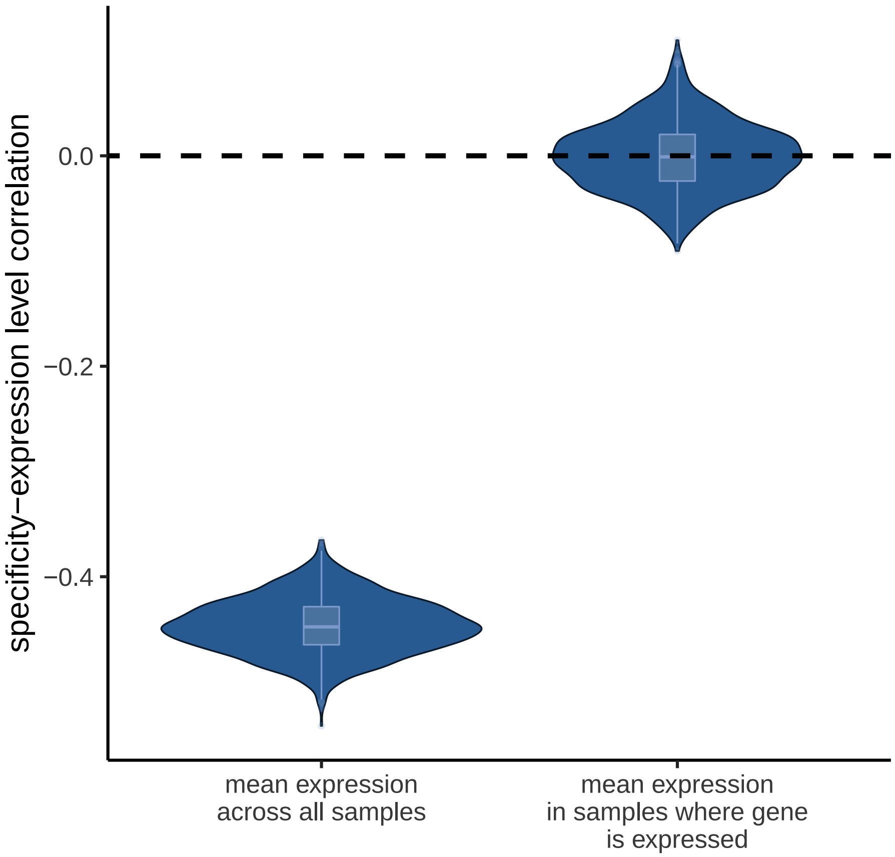


Figure S39: Violin plot of correlations between expression specificity and expression level across 1000 simulated expression matrices. In the two different simulations shown here, expression level was quantified either as average expression across all samples or average expression across samples with expression values $> 0$. Each expression matrix was simulated by sampling from a zero-inflated negative binomial distribution.
